# Machine Learning Resolves Functional Phenotypes and Therapeutic Responses in KCNQ2 Developmental Epileptic Encephalopathy iPSC Models

**DOI:** 10.1101/2025.07.26.666977

**Authors:** Dina Simkin, Syed M.A. Wafa, Mennat Gharib, Kelly A. Marshall, Yezi Yang, Linda C. Laux, Alfred L. George, Evangelos Kiskinis

## Abstract

Pathogenic KCNQ2 variants are associated with developmental and epileptic encephalopathy (KCNQ2-DEE), a devastating disorder characterized by neonatal-onset seizures and impaired neurodevelopment with no effective treatments. KCNQ2 encodes the voltage-gated potassium channel K_V_7.2, which regulates action potential threshold and repolarization. However, the relationship between K_V_7.2 dysfunction and abnormal neuronal activity remains unclear. Here, we use human induced pluripotent stem (iPSC)-derived neurons from 5 KCNQ2-DEE patients with pathogenic variants and CRISPR/Cas9-corrected isogenic controls to investigate pathophysiological mechanisms. We identify a common dyshomeostatic enhancement of Ca^2+^-activated small conductance potassium (SK) channels, which drives larger post-burst afterhyperpolarizations in KCNQ2-DEE neurons. Using microelectrode arrays (MEAs), we recorded over 18 million extracellular spikes from >8,000 neurons during 5 weeks in culture and then applied supervised and unsupervised machine learning algorithms to dissect time-dependent functional neuronal phenotypes that defined both patient-specific and shared firing features among KCNQ2-DEE patients. Our analysis identified irregular spike timing and enhanced bursting as functional biomarkers of KCNQ2-DEE and demonstrated the significant influence of genetic background on phenotypic diversity. Importantly, using unbiased machine learning models, we showed that chronic treatment with the K_V_7 activator retigabine rescues the disease-associated functional phenotypes with variable efficacy. Our findings highlight SK channel upregulation as a critical pathophysiological mechanism underlying KCNQ2-DEE and provide a robust MEA-based machine learning platform useful for deciphering phenotypic diversity amongst patients, discovering functional disease biomarkers, and evaluating precision medicine interventions in personalized iPSC neuronal models.

## Introduction

Heterozygous pathogenic variants in KCNQ2 encoding the voltage-gated K^+^ channel K_V_7.2 are the leading cause of neonatal-onset epilepsy^1-3^. KCNQ2-related disorders present along a wide clinical spectrum, ranging from self-limited familial neonatal epilepsy (SLFNE) featuring neonatal-onset seizures abating in late infancy followed by normal neurodevelopment, to severe developmental and epileptic encephalopathy (DEE) characterized by neonatal-onset seizures and lifelong cognitive, developmental, and motor deficits^4-7^. The factors driving diverse clinical presentations are unclear. KCNQ2-encoded channel subunits co-assemble with related K_V_7.3 to form heterotetrameric channels that generate the neuronal M-current^8,9^, a slowly activating, non-inactivating, voltage-dependent K^+^ conductance. M-current regulates neuronal excitability by setting the action potential (AP) threshold and contributing to AP repolarization as well as post-burst afterhyperpolarization (AHP).

Although studies in heterologous expression systems have determined that most KCNQ2-DEE mutations exhibit loss-of-function often with dominant-negative effects, there is not always a clear distinction in biophysical deficits between variants associated with SLFNE and DEE^10^. Heterologous models do not replicate the native neuronal context, potentially missing expression-level deficits such as altered stability and mislocalization of mutant channels, as well as any potential compensatory responses of developing neurons. Additionally, identical KCNQ2 mutations can yield divergent clinical phenotypess^11^, suggesting that channel properties alone do not fully predict their impact and highlighting the need to understand contributions of genetic background on disease outcome.

Several investigative studies have employed mouse models with KCNQ2 knockouts^12-18^ or disease-related KCNQ2 missense mutations^18-31^. These models have invariably demonstrated the association between KCNQ2 channel dysfunction, seizures, and behavioral phenotypes in mice. However, key developmental differences restrict the translational value of mouse models^32-49^, as seizures in KCNQ2-DEE patients manifest within days of birth during a crucial period of human cortical expansion and synaptogenesis, whereas seizures in mouse models typically emerge at later stages^22,23,31,50^ corresponding to human adolescence^51^. Additionally, mouse studies have been conducted independently, with varying genetic backgrounds and different approaches for introducing mutations, making it challenging to discern whether identified phenotypes are unique to specific mutations or indicative of shared pathogenic pathways underlying the broader spectrum of KCNQ2-related disorders. These limitations underscore the value of patient-specific induced pluripotent stem cell (iPSC)-derived neuronal models that allow the examination of KCNQ2 mutations in human neurons, and in each patient’s unique genetic background, to better define shared and patient-specific disease mechanisms. We previously demonstrated that KCNQ2-DEE cortical excitatory neurons, from a single patient carrying a heterozygous KCNQ2 loss-of-function mutation (R581Q), exhibited a progressive shift in their functional phenotype that was first consistent with a loss of M-current, but later became dependent on dyshomeostatic upregulation of other K^+^ channels^52^. In this study, we examined the molecular and functional phenotype heterogeneity in neurons across an expanded set of 5 KCNQ2-DEE patient-derived and paired isogenic control iPSC lines to define common disease mechanisms and establish a platform that can be used to identify patient-specific therapeutic responses.

## Results

### Generation of multiple pairs of patient-specific KCNQ2-DEE and mutation-corrected isogenic control iPSC lines and neurons

We established a cohort of 5 KCNQ2-DEE iPSC lines generated from patients recruited at Ann & Robert H. Lurie Children’s Hospital of Chicago. All 5 individuals were heterozygous for distinct missense *KCNQ2* variants (R207W, H228R, T274M, P335L, R581Q), exhibited seizures during the first few days after birth, and had severe developmental delay (Figure 1A-B). Importantly, all variants were shown to diminish KCNQ2 current density or alter its activation voltage-dependence to varying degrees in heterologous expression models^10^. The iPSC lines were produced by reprogramming peripheral blood mononuclear cells (PBMCs) and all mutations were corrected by CRISPR/Cas9 gene editing to generate isogenic control lines (Figure 1C). We performed quality control for genomic integrity and pluripotency using a combination of assays including targeted Sanger sequencing at the site of the mutation, karyotype analysis, staining with key pluripotency markers, and allele copy number genomic qPCR analysis (Figure S1A-C)^53^. All patient-specific and isogenic control iPSC lines were systematically differentiated into cortical excitatory glutamatergic neurons using the well-established *Ngn2* transcription factor-based induction protocol^52^ (Figure 1C-D). The efficiency of differentiation among lines was quantified using immunocytochemistry (ICC) with neuronal markers MAP2 and vGLUT1 (Figures 1D and S1D) to find that ≥79% of neurons were GFP positive (Figure S1E), as well as ≥99% and ≥98% MAP2 and vGLUT1 positive, respectively (Figure S1F-G).

**Figure 1.**
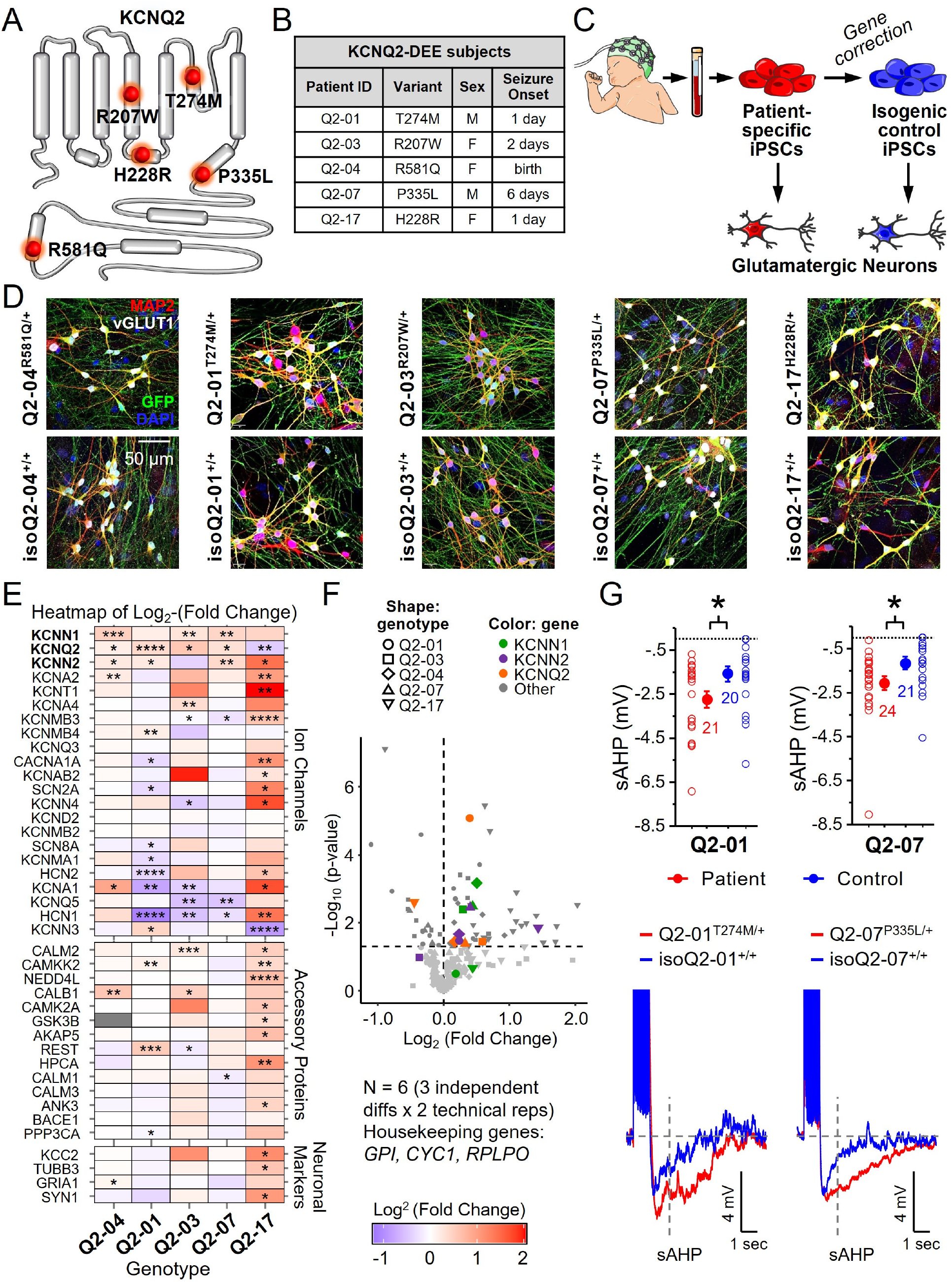
Cortical excitatory neurons from different KCNQ2-DEE patient-specific iPSC lines exhibit a converging cellular phenotype. **A)** Illustration of proposed structure of KCNQ2 channel subunit highlighting the locations of KCNQ2-DEE mutations investigated in this study: R207W at the S4 voltage-sensing domain, H228R at the intracellular S4-S5 linker, T274M at the pore domain, and P335L and R581Q at the C-terminus. **B)** Table summarizing molecular and clinical characteristics of the participants in this study, including subject ID, KCNQ2 variant, diagnosis sex, and age at seizure onset. **C)** Schematic of the iPSC-based platform used to generate cortical excitatory neurons. Image adapted from Servier Medical Art (https://smart.servier.com/), licensed under CC BY 4.0. **D)** Immunocytochemical labeling of paired KCNQ2-DEE and isogenic control neurons at day 26, showing glutamatergic (vGLUT1) and neuronal (MAP2) markers, along with GFP and DAPI. Scale bar: 50 µm. **E)** Heatmap of day 35 neuron gene expression profiles across 5 KCNQ2-DEE patient iPSC lines (log_2_ fold-change relative to respective isogenic controls; N = 6 (3 independent differentiations x 2 technical replicates per line). Forty-two gene targets are first grouped by class (ion channels, accessory proteins, neuronal markers) and ranked within each class by the average log_2_ (p-values) x sign of fold change, showing significant fold-change up-regulation (top) or down-regulation (bottom). Red = up-regulated, purple = down-regulated; grey = not tested. Housekeeping genes: GPI, CYC1, RPLPO. t-test: *p < 0.05, **p < 0.005, ***p < 0.0005 **F)** Volcano plot of gene expression data on day 35 displaying -log10(p-value) plotted against log2(fold-change). Neurons from different KCNQ2-DEE lines are represented using different shapes. KCNN1, KCNN2, and KCNQ2 are highlighted in distinct colors to indicate robust and consistent up-regulation across patients. Horizontal dashed line marks the 5% significance threshold; vertical dashed line indicates ±1-fold (log_2_ = ±1). **G)** Whole-cell current-clamp recordings were used to measure the slow post-burst AHP (sAHP) following a 50 Hz train of 25 action potentials evoked by 2 ms/1.2 nA suprathreshold current injection steps in patient and isogenic control neurons from Q2-01 and Q2-07 pairs of lines during day 22-26. Top: KCNQ2-DEE (red) neurons exhibit larger sAHP compared to respective isogenic controls (blue; t-test: Q2-01, p = 0.0163; Q2-07, p = 0.0202). Bottom: representative voltage traces from Q2-01 and Q2-07 lines showing post-burst AHPs measured 1 second after the end of stimulus. The number of neurons analyzed per group is displayed within the figure and in Table S1. Data are shown as mean ± SEM.

### KCNQ2-DEE patient iPSC neurons exhibit upregulation of SK channel expression and enhanced slow post-burst afterhyperpolarizations (AHPs)

To investigate time-dependent disease mechanisms in this cohort of patient-specific neurons, we interrogated the mRNA expression of 40 genes involved in the regulation of neuronal excitability, including ion channels (n=22), ion channel accessory proteins (n=14), and neuronal markers (n=4). We performed RT-qPCR at early (day 15) and late (day 35) time points following *in vitro* neuronal differentiation (Figures 1E and S2). Gene expression levels from three independent neuronal differentiations for each pair of lines (e.g., KCNQ2-DEE and isogenic control) were first normalized to the average level of three housekeeping genes and subsequently, each patient was normalized to the respective mutation-corrected control. While there was significant diversity in gene expression profiles, we found consistent upregulation of the SK channel genes *KCNN1* and *KCNN2* as a shared feature across all KCNQ2-DEE patient iPSC neurons at day 35, with *KCNN2* significantly upregulated in 4 out of 5 cases, and *KCNN1* in 3 out of 5 cases (Figure 1E-F). We also found upregulation of *KCNQ2* itself, which may reflect a direct compensatory mechanism for the underlying KCNQ2 loss-of-function. Critically, besides Q2-17^H228R/+^, which exhibited increased expression of *KCNN1* and *KCNN2* at day 14, these alterations emerged only at the later time point (Figure 1E and S2).

The upregulation of the two small-conductance Ca^2+^-activated K^+^ (SK) channels can translate into electrophysiological changes, such as enhanced post-burst AHPs in neurons. To investigate this possibility, we performed current-clamp recordings to compare the intrinsic excitability of Q2-01^T274M/+^ and Q2-07^P335L/+^ neurons to their respective isogenic controls on days 22-26 in culture (Figure 1G and Table S1). While we did not detect substantial differences in the passive or action potential waveform properties between KCNQ2-DEE and isogenic control neurons (Table S1), we found that the slow Ca^2+^- dependent post-burst AHPs were enhanced in both Q2-01^T274M/+^ and Q2-07^P335L/+^ neurons (Figure 1G). Collectively, these results are consistent with our prior findings in Q2-04^R581Q/+^ neurons^52^, and suggest that SK channel upregulation and the associated enhancement of post-burst AHPs may represent a shared pathological feature of KCNQ2-DEE neurons, highlighting the maladaptive nature of this compensatory mechanism in regulating excitability.

### KCNQ2-DEE patient iPSC neurons exhibit common, time-dependent spontaneous functional phenotypes on microelectrode arrays (MEAs)

To investigate the functional ramifications of the transcriptional changes we identified, we used 48-well MEA plates to record extracellular spikes daily from day 9 to 35 across all 5 KCNQ2-DEE and isogenic control neuron pairs (Figure 2A-B). Notably, we observed that as neurons matured and began to exhibit robust and consistent activity (Figure 2B), mere media changes had pronounced effects on neuronal firing across all cell lines (Figure S3A). To quantify this observation, we decomposed firing rates into “trend” and “seasonal” components (i.e., regularly recurring fluctuations in firing tied to media changes), which respectively capture long-term shifts and cyclical patterns in time-series data (Figure S3B). Periodic fluctuations in the seasonal component emerged around day 19 in both pooled KCNQ2-DEE neurons and pooled isogenic controls, with no observable differences between them (Figure S3C-G). The moderate strength of the seasonal component (0.53-0.54; Figures S3D and S3H), reflecting the degree to which neuronal activity cycles aligned with feeding schedule, established media changes as a confounding variable during recordings (Figures S3E and S3I), with its influence increasing sigmoidally over time (Figures S3F and S3J). Interestingly, we observed deeper troughs on non-feeding days compared to shallow peaks on feeding days, suggesting that firing suppression on non-feeding days was stronger than firing enhancement on feeding days (Figures S3D and S3H). Thus, to maximize data inclusion during early developmental periods whilst mitigating later confounding effects, we sampled every recording until day 16, after which only recordings from feeding days were used for subsequent analysis.

**Figure 2.**
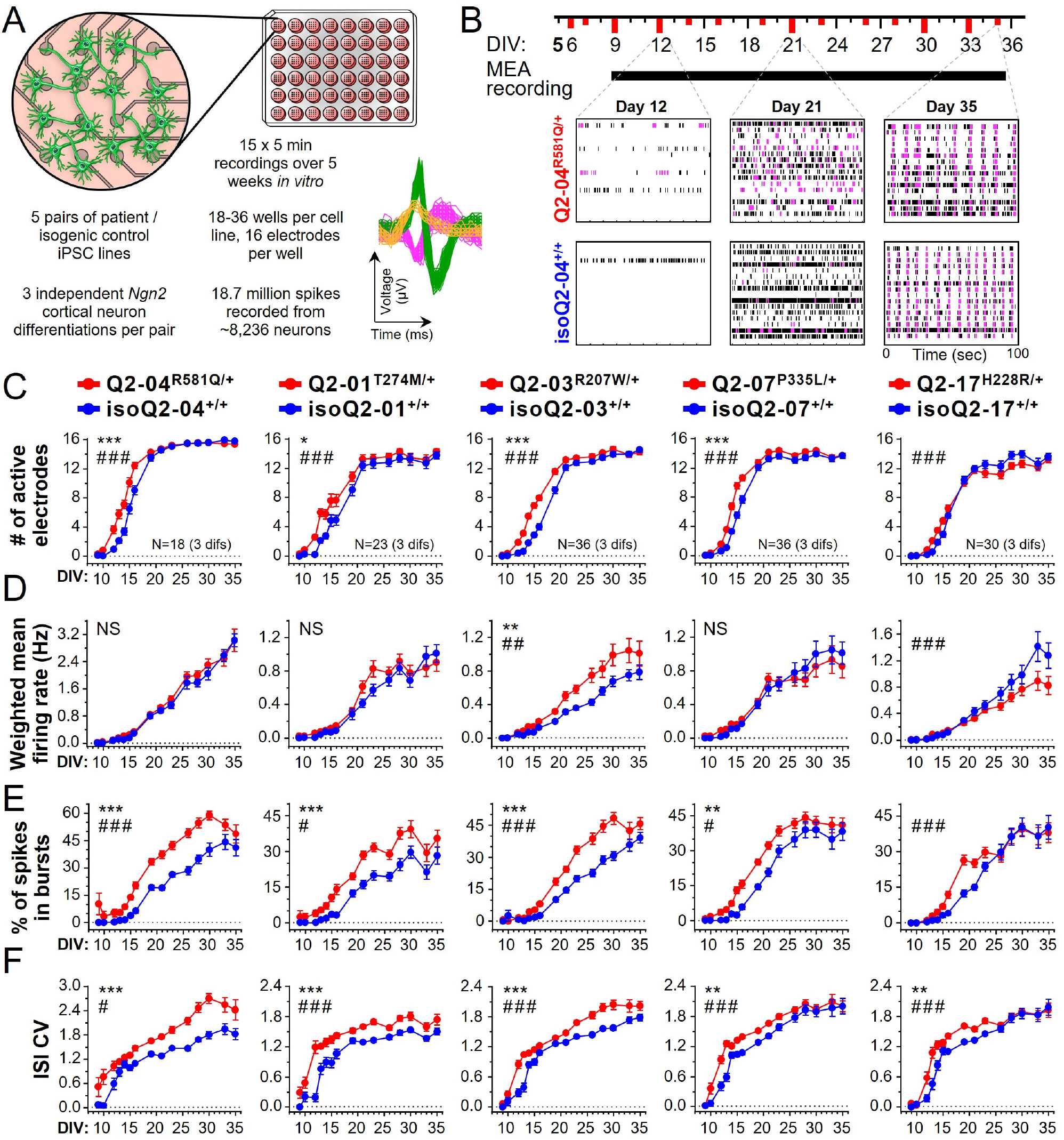
MEA analysis reveals early onset of spontaneous activity and enhanced bursting propensity across cortical excitatory neurons derived from 5 KCNQ2-DEE patient iPSC lines. **A)** Illustration of a 48-well MEA plate with 16 electrodes per well. Data were collected from 5 pairs of KCNQ2-DEE and isogenic control neurons with 3 independent cortical neurons differentiations per pair. Spontaneous activity of neurons was recorded in 15 five-minute sessions during 5 weeks in culture. **B)** Illustration of the recording time course, with neurons plated on day 5. Red ticks denote days when half of the neuronal culture media was replaced with fresh media. Bottom: Spike-activity raster plots showing spontaneous activity across 100 seconds from a single representative MEA well of KCNQ2-DEE (Q2-04R581Q/+) and isogenic control (isoQ2-04+/+) neurons at days 12, 21, and 35. Each row represents one of 16 electrodes; black ticks indicate single spikes, and pink ticks denote spikes occurring within bursts. **C)** Longitudinal analysis of the number of active electrodes per well. Across all 5 pairs, KCNQ2-DEE neuron cultures recruited a larger fraction of electrodes during the first ∼10 days of spontaneous firing (days 9-19) relative to isogenic controls, indicating earlier onset of activity (repeated measures ANOVA: *genotype effect; #genotype/time interaction). Differences were not observed at later time points (day ≥ 21). Data are shown as mean ± SEM, and the number of replicate wells is indicated in the panel. **D)** Longitudinal analysis of weighted mean firing rate (WMFR) showed no consistent effects across KCNQ2-DEE patient iPSC lines. However, there were line-specific differences with elevated WMFR in Q2-03R207W/+ and reduced in Q2-17H228R/+ neurons, with no significant effect (NS) in genotype or interaction of genotype/time in the remaining three lines (repeated measures ANOVA). **E)** Longitudinal analysis of the percentage of all spikes occurring within bursts (burst %) revealed that all KCNQ2-DEE patient iPSC neurons fired a significantly greater fraction of their spikes within bursts, suggesting enhanced bursting activity (repeated measures ANOVA). **F)** Longitudinal analysis of inter-spike interval coefficient of variation (ISI CV) revealed that KCNQ2-DEE patient iPSC neurons exhibited higher ISI CV throughout the recording period, indicating increased spike time irregularity, which is indicative of enhanced bursting. Repeated-measures ANOVA: genotype effect: *p < 0.05, **p < 0.005, ***p < 0.0005; genotype/time interaction: # p < 0.05, ## p < 0.005, ### p < 0.0005. Data are shown as mean ± SEM; n = 18-36 technical replicates (wells) per cell line from three independent differentiations.

All KCNQ2-DEE neurons exhibited greater numbers of active electrodes between day*s* 9 and 19 compared to their respective isogenic controls, indicating an earlier onset of neuronal activity (Figure 2C). Because there were no differences in the number of active electrodes at later time points, this early activity was not attributed to differences in neuronal density. Nevertheless, to determine whether different numbers of recorded neurons could confound downstream measures of excitability^54^, we performed unsupervised spike sorting^55,56^ on spike waveforms on the last day of recording (day 35). Spike waveforms recorded from each electrode were reduced to a few dimensions and the optimal number of distinct waveforms was determined by clustering the resultant embeddings (Figure S4A), which we used as a proxy for the number of active neuronal units. On average, spikes recorded from each electrode were sorted into approximately 2 groups (1.83 ± 0.03 distinct clusters/electrode), suggesting that each electrode recorded electrical activity from ∼2 neurons. Importantly, there were no differences in the number of estimated neurons between patient and paired isogenic control cultures, suggesting that earlier onset of activity is an intrinsic developmental property of KCNQ2-DEE neurons (Figure S4B).

To determine the impact of *KCNQ2* mutations on neuronal activity, we quantified several single electrode metrics across time in culture. While neurons from three of the KCNQ2-DEE patient iPSC lines did not exhibit significant alterations in the basic weighted mean firing rate (WMFR), two lines exhibited divergent excitability, with Q2-03^R207W/+^ neurons showing an enhancement, and Q2-17^H228R/+^ a slight reduction relative to their respective isogenic controls (Figure 2D). In contrast, neurons from all 5 KCNQ2-DEE lines exhibited a strong propensity for firing in bursts as indicated by activity raster plots (Figure S5A). Quantifying this observation, we found that all KCNQ2-DEE neurons consistently exhibited a higher percentage of spikes occurring in bursts (Burst %; Figure 2E) and larger interspike interval coefficient of variation (ISI CV; Figure 2F), reflecting greater irregularity in spike timing associated with enhanced bursting activity. While irregular spiking and enhanced bursting are consistent with SK channel upregulation^57^, these differences diminished toward the end of the recording time course, suggesting that other functional features may drive disease mechanisms later in development. These findings demonstrate the time-dependent nature of functional activity phenotypes in neurons harboring pathogenic *KCNQ2* mutations. At the same time, these findings highlight the complexity of pathophysiological mechanisms underlying KCNQ2-DEE and emphasize the need for rigorous and unbiased computational methods to analyze multi-parametric longitudinal recordings.

### A comprehensive computational framework reveals time-dependent functional profiles of KCNQ2-DEE patient iPSC neurons

Our dataset of MEA recordings represents a unique resource comprising 18 million extracellular spikes recorded from multiple KCNQ2-DEE patient and isogenic control neuron pairs during five weeks in culture (n=3 independent differentiations). From these recordings, we curated a set of 41 distinct functional features that collectively capture the complexity of population-level neuronal activity. However, analyzing individual features in isolation risks overlooking the broader, time-dependent, multi-parametric nature of the KCNQ2-DEE disease phenotype. To interrogate this complex dataset in an unbiased and scalable way, we established an integrated computational strategy. Given the longitudinal sparsity of our recordings and the presence of systematic batch effects^58,59^, we developed a comprehensive, statistically rigorous computational framework to process, analyze, and visualize all 41 activity-based features (Figure 3). Briefly, our computational framework consisted of three steps to address the technical limitations of functional phenotyping assays (Figure 3A; See methods). First, we resampled features using linear interpolation to account for exclusion of recordings conducted at non-feeding timepoints (Figure S3). Notably, although interpolated data were used for downstream analyses, they were excluded from statistical hypothesis testing. Second, we scaled features to account for batch effects and make them comparable to one another. Third, we subtracted mean scaled feature values of isogenic controls at each timepoint from scaled feature values of each culture well, allowing effect sizes to be compared consistently across time. Collectively, these steps allowed us to visualize the temporal and multiparametric functional profiles of individual KCNQ2-DEE neurons and revealed both shared and patient-specific firing patterns (Figure 3B). To enable accessibility and utilization of this MEA data analysis tool, we created a freely available web-based application (kiskinislab.shinyapps.io/mea-snap/).

**Figure 3.**
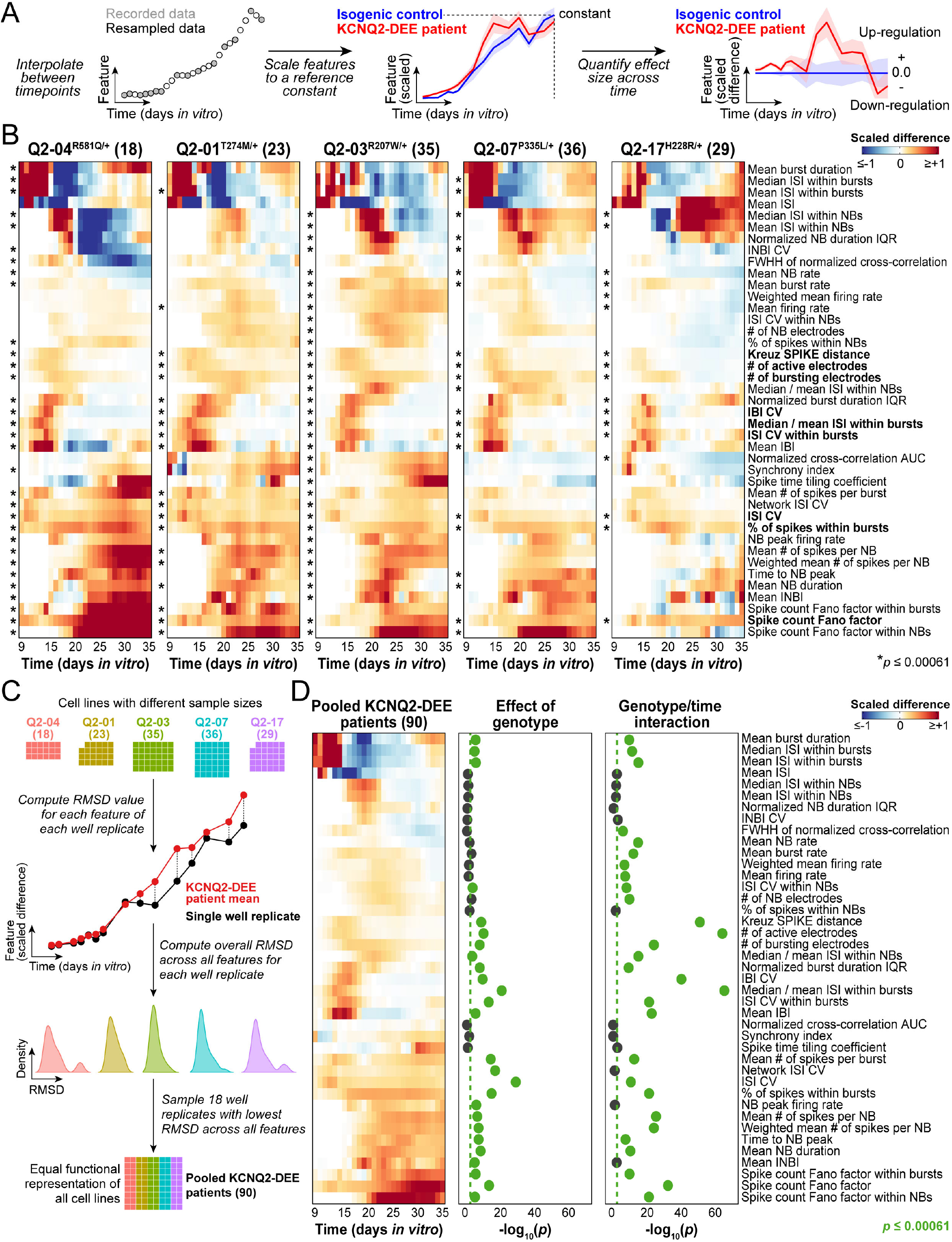
A computational framework resolves time-dependent functional profiles of patient-specific KCNQ2-DEE iPSC neurons. **A)** Schematic of the MEA data processing workflow. Each of the 41 extracted functional features underwent a three-step signal processing workflow: (1) linear interpolation to address exclusion of non-feeding day recordings, (2) feature scaling to normalize values and enable comparison across features, batches, and patient-control pairs, and (3) scaled differencing to quantify genotype-associated effect sizes over time (see methods for details). **B)** Heatmaps showing scaled differences in functional features between KCNQ2-DEE neurons and their respective isogenic controls. Positive values indicate up-regulation in patient-derived neurons, while negative values indicate down-regulation. \**p* ≤ 0.00061 based on repeated measures ANOVA assessing genotype or interaction between genotype and time. **C)** To account for different sample sizes among patient-derived iPSC lines, root mean square deviation (RMSD) values were computed between each culture well (technical replicate) and the mean of the cell line. Equal numbers of wells with the lowest RMSD values were selected, ensuring equal representation of all patient-derived cell lines for downstream analysis. **D)** Heatmap summarizing scaled differences in functional features between pooled KCNQ2-DEE neurons and pooled isogenic controls. Corresponding log_10_(*p*-values) of genotype and interaction between genotype and time are shown. Green dashed vertical lines represent the significance threshold (p = 0.05), and green circles indicate statistically significant effects (*p* ≤ 0.00061).

Statistical hypothesis testing on the processed high-dimensional dataset identified nine functional features that showed significant genotype effects and/or interactions between genotype and time across all KCNQ2-DEE patient lines (repeated measures ANOVA, *p* ≤ 0.00061 after Bonferroni correction for multiple comparisons). These features fell into two broad, time-dependent categories. Early in the neurodevelopmental time course, KCNQ2-DEE neurons exhibited a larger number of active electrodes and higher Kreuz SPIKE-distance synchrony compared to their respective isogenic controls, indicating an earlier onset of coordinated neuronal activity (Figures 3B and S6). This early activity was accompanied by more variable bursting behavior, with greater numbers of bursting electrodes, greater ISI CV within bursts, higher median/mean inter-spike interval (ISI) within bursts ratios, and greater inter-burst interval CV (Figures 3B and S6). The simultaneous rise in the median/mean ISI ratio and ISI CV within bursts suggests a reduction in spike time accommodation but greater overall spike time variability within bursts, indicating more irregular and/or unstable burst structure. As development progressed, this initial divergence was followed by more variable spike timing and higher bursting propensity, evidenced by elevated spike count Fano factors, greater ISI CV, and greater percentage of spikes occurring within bursts (Figure 3B and S7). Together, these findings demonstrate that KCNQ2-DEE neurons harboring distinct heterozygous mutations exhibit common electrophysiological abnormalities, marked by greater variability in spike timing and bursting patterns, and suggest shared functional sequelae across patients.

To determine the average functional profile of KCNQ2-DEE neurons, we first down-sampled replicate wells so that neurons from all 5 patient lines were equally represented. We subsequently used root mean square deviation (RMSD) values to select representative culture wells based on their similarity to group means across all features (Figure 3C). Pooled KCNQ2-DEE neurons exhibited higher numbers of active populations that fired earlier and more synchronously, followed by bursting abnormalities and higher firing irregularity, relative to mutation-corrected isogenic controls (Figure 3D). Statistical analysis across this aggregated dataset identified 32 functional features with significant genotype effects and/or interactions between genotype and time (Figure 3D), likely reflecting greater statistical power with the large sample size.

### Supervised machine learning discovers developmental stages and functional biomarkers of KCNQ2-DEE patient iPSC neurons

Because statistically significant features may not all be equally relevant for defining the disease biologically, we next developed a supervised machine learning framework to identify the most predictive functional biomarkers of KCNQ2-DEE neurons. Inspired by prior work that used functional features to predict the maturation of primary rat cortical cultures^60^, we trained supervised machine learning classifiers to distinguish pooled KCNQ2-DEE neurons from pooled isogenic controls. By quantifying feature contributions to classification performance over time, we identified time-dependent functional biomarkers of KCNQ2-DEE. Briefly, we trained machine learning algorithms known as “ensembles of decision trees” at each time point to classify culture wells as either KCNQ2-DEE or control (Figure 4A)^61^. Unlike black-box artificial intelligence models, decision trees are interpretable algorithms that make a series of binary decisions based on individual features, progressively partitioning the feature space to create clear class boundaries that maximize separation between the two classes^62-64^ (Figure 4B). The inherent interpretability of decision trees makes them especially valuable in pre-clinical models, where understanding feature importance is valuable for biomarker discovery. To ensure that machine learning classifiers were trained and tested on balanced distributions of biological replicates and classes, we randomly assigned equal proportions of each KCNQ2-DEE variant and its corresponding isogenic control to training (16 replicate wells per cell line; *n* = 160, 89% of dataset) or test (2 replicate wells per cell line; *n* = 20, 11% of the dataset) sets (Figure 4C). To improve the ability of classifiers to generalize to new and unseen data, we further split the training set into four parts and implemented 4-fold cross-validation, where classifiers are trained on three parts (*n* = 120, 75% of the training dataset) then validated on the remaining part (*n* = 40, 25% of the training dataset) in a rotating fashion. During this process, we also fine-tuned hyperparameters. Independent classifiers were trained on all 41 firing features from each day in culture, and feature importance was quantified by randomly shuffling feature values and measuring resulting drops in classifier accuracies (Figure 4C). The accuracy of the classifier in identifying KCNQ2-DEE neurons measured by the cross-validation F_1_-score, started at 67.9% on day 9, peaked at 89.5% on day 13 and remained relatively stable through day 22, before declining to 74.1% by day 35 (Figure 4D, top).

**Figure 4.**
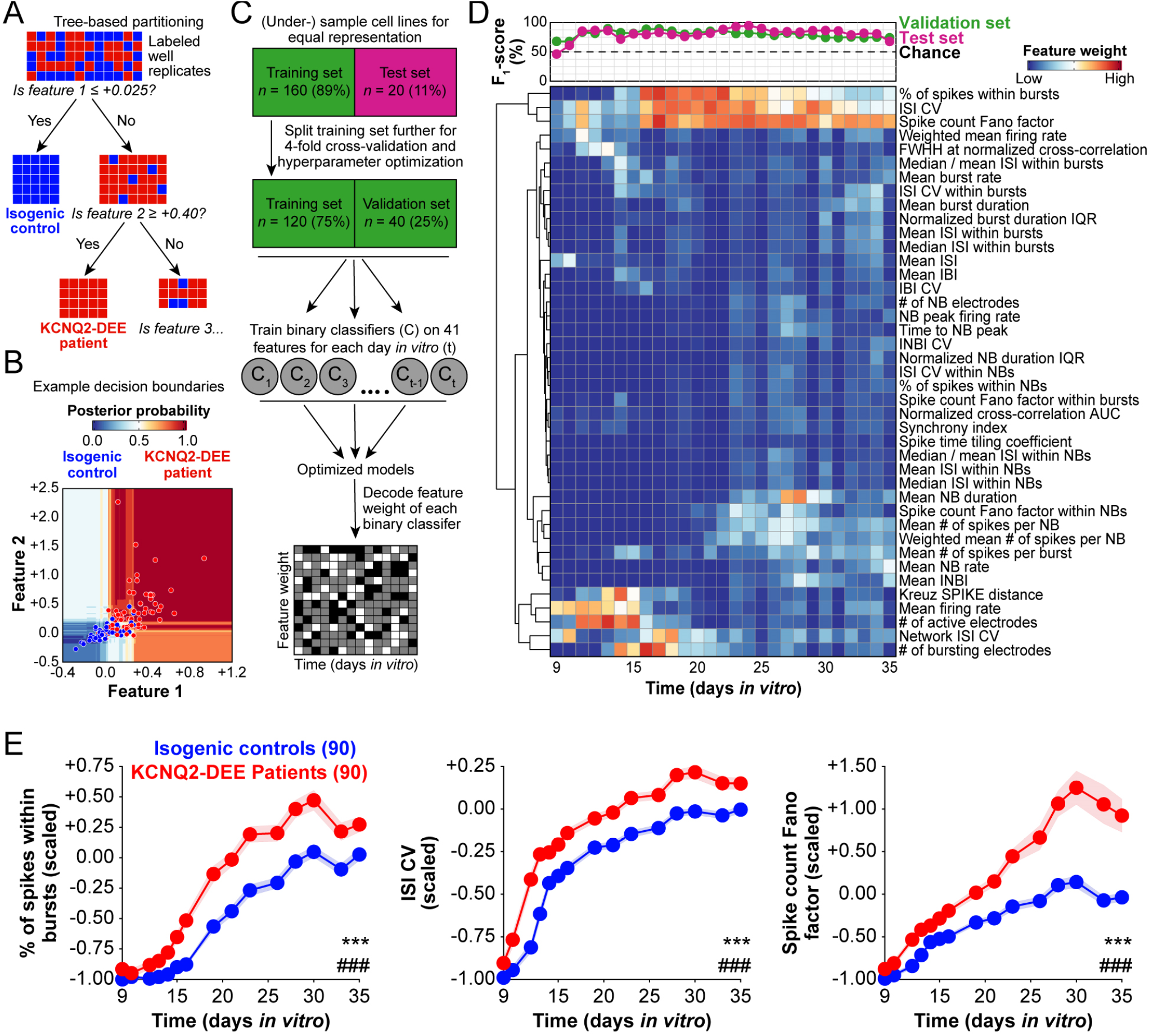
Supervised machine learning enables identification of time-dependent functional biomarkers of KCNQ2-DEE. **A)** Schematic illustrating how tree-based classifiers recursively split data by posing a series of binary (‘yes’ or ‘no’) questions based on feature thresholds to separate samples into classes. **B)** Example decision boundaries from a binary classifier trained on two features, demonstrating how tree-based models segment the feature space into regions associated with predicted class membership. The background shading reflects posterior probabilities, with darker shades indicating greater model confidence in class assignment. **C)** Workflow for supervised classification, showing the splitting of the dataset into training, cross-validation, and test sets for model training, hyperparameter optimization, performance evaluation, and feature importance estimation. **D)** Heatmap showing the relative normalized importance of each feature in distinguishing KCNQ2-DEE neurons from controls across time. Higher values indicate a greater contribution of the feature to classification. Classifier performance on cross-validation and held-out test sets are indicated above the heatmap. **E)** The top three most predictive functional biomarkers of KCNQ2-DEE. Compared to control neurons, KCNQ2-DEE neurons exhibit a higher percentage of spikes within bursts, higher ISI CV, and higher spike count Fano factor indicating greater bursting propensity and greater variability in spike timing. Repeated-measures ANOVA: genotype effect: *p < 0.05, **p < 0.005, ***p < 0.0005; genotype/time interaction: # p < 0.05, ## p < 0.005, ### p < 0.0005. Data are shown as mean ± SEM; and the number of replicate wells is indicated in the panel.

The accuracy of the model was driven by several functional neuronal features. Time-resolved analysis of classifier weights revealed three firing features that exhibited a similar temporal pattern and a strong predictive value of KCNQ2-DEE neurons (Figure 4D-E and S8). These included the percent (%) of spikes within bursts, the ISI CV, and spike count Fano factor (Figure 4D-E). Notably, spike count Fano factor, which measures the regularity of neuronal spike timing, emerged as the most consistently predictive functional biomarker of KCNQ2-DEE neurons across time, suggesting its potential use as a pre-clinical endpoint for assessing candidate therapeutics (Figure 4D-E). Further analysis of model weights revealed two overlapping stages during the *in vitro* development of KCNQ2-DEE neurons (Figure 4D). In stage one (days 9-19), early changes in onset of activity, excitability, synchrony, and bursting were most predictive of KCNQ2-DEE neurons. Stage two (days 16-35) featured shifts in bursting population dynamics and spike timing regularity, with pooled KCNQ2-DEE neurons exhibiting enhanced bursting propensity and more irregular firing patterns. Interestingly, the predictive value of bursting propensity gradually declined after day 26, while disruptions in firing regularity persisted through the end of the recording period on day 35. One possible explanation is that, as activity shifts toward coordinated network bursts over time, spikes become more tightly clustered within network bursts. Because network-burst duration displays higher predictive weight in our machine-learning model (days 26-28; Figure 4D and S8), metrics tuned to short, well-defined bursts like % of spikes in bursts and ISI CV may lose discriminative power. On the other hand, the spike count Fano factor may remain elevated because extended network bursts still produce irregular spike timing and variable spike counts across time bins.

### Unsupervised machine learning demonstrates the strong influence of genetic background and KCNQ2 mutations on the functional phenotype of neurons

The identification of different developmental stages of firing patterns (Figure 4), as well as the significant differences of firing features based on genotype over time interactions (Figure 2), suggests that *KCNQ2* mutations impact the functional trajectory of iPSC-derived neurons. To quantify functional trajectories across time and uncover any potential KCNQ2-dependent alterations, we transformed the MEA recording dataset into a lower-dimensional space and performed cluster analysis to uncover spatiotemporal patterns. Consistent with previous studies^58^, we found that standard dimension reduction techniques including principal component analysis (PCA), *t*-distributed stochastic neighbor embedding (t-SNE), and unified manifold approximation and projection (UMPA), failed to capture the temporal dynamics in our dataset (Figure S9A, see Methods). In contrast, temporal potential of heat diffusion for affinity-based transition embedding (T-PHATE)^65^ successfully recovered a low-dimensional intrinsic manifold structure from our high-dimensional electrophysiological feature dataset, and separated neurons based on longitudinal functional dynamics along days 9-35 *in vitro* (Figures 5A and S9A). T-PHATE achieves this by explicitly incorporating temporal autocorrelation, capturing how neuronal activity at one time point relates to adjacent time points. Projecting each KCNQ2-DEE and isogenic control profile onto the lower-dimensional latent structure revealed distinct functional trajectories amongst individuals, driven primarily by genetic background (Figure 5B and S9B).

**Figure 5.**
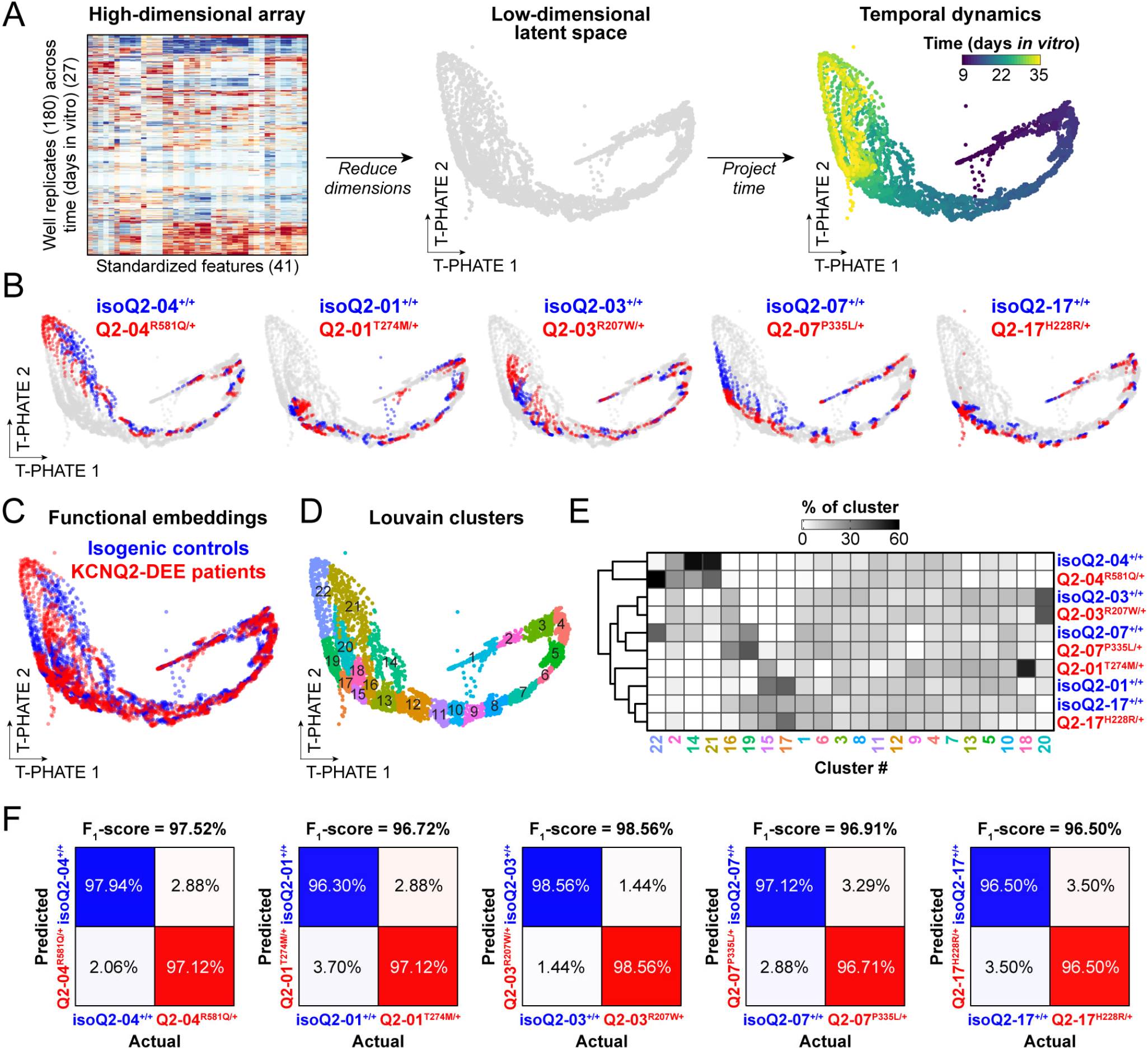
Dimensionality reduction using T-PHATE reveals the influence of genetic background on the functional trajectories of patient-specific KCNQ2-DEE iPSC neurons. **A)** Schematic illustrating the embedding of neuronal cultures from a high (41)-dimensional feature space into a low (3)-dimensional latent space across time using T-PHATE. Projecting time onto the lower-dimensional structure reveals smooth trajectories that capture the temporal dynamics of neuronal activity. **B)** Overlaying individual cell line labels onto the latent space demonstrates that functional variation is strongly structured by genetic background. **C)** Overlay of all KCNQ2-DEE neurons and isogenic controls within the latent T-PHATE space. **D)** A total of 22 clusters were identified using the Louvain algorithm, which detects communities based on local neighborhood connectivity in the manifold. **E)** Heatmap showing the percentage contribution of each iPSC-derived neuronal cell line to each cluster. The accompanying dendrogram further supports the influence of genetic background in organizing phenotypic diversity. Among the 5 genetic backgrounds, Q2-01 and Q2-17 cell lines exhibited the most similar functional trajectories, followed by Q2-03 and Q2-07. In contrast, neurons derived from the Q2-04 background were the most functionally divergent. **F)** Supervised classifiers trained to distinguish KCNQ2-DEE neurons from their respective isogenic controls using T-PHATE embeddings achieve high cross-validation accuracies (≥96.50%) suggesting that, within each genetic background, KCNQ2-DEE neurons and isogenic controls follow distinct functional trajectories.

To quantify the spatiotemporal and functional relationships among patient iPSC neurons, we next used Louvian clustering, to find 22 clusters of neurons across space and time defined by their electrophysiological features (Figures 5C-D). Hierarchical clustering and calculation of the percentage of each neuron cluster occupied by each genotype captured the strong influence of genetic background, as lines separated based on patient genotype and not based on *KCNQ2* genotype (Figure 5E). These findings are in accordance with prior studies that have highlighted the importance of genetic background on neuronal differentiation propensity and maturation^66,67^, although our analysis provides the first direct evidence that genetic background significantly shapes the electrophysiological trajectories of neurons. Importantly, while we found a dominant effect of genetic background on driving the functional trajectory of iPSC-derived neurons, the impact of *KCNQ2* mutations was also visually evident, as KCNQ2-DEE neurons (red) appeared distinct from their paired mutation-corrected isogenic control neurons (blue; Figure 5B and S9B). To investigate this observation further, we trained supervised machine learning classifiers to distinguish T-PHATE functional embeddings of KCNQ2-DEE genotypes within each genetic background. The classifiers were able to successfully distinguish KCNQ2-DEE neurons from their corresponding isogenic control neurons control neurons with high accuracy (≥96.50%; Figure 5F), demonstrating that KCNQ2 channel dysfunction significantly alters the functional trajectory of iPSC-derived excitatory neurons.

### Machine learning predictions and gene expression analyses demonstrate that KV7 channel inhibition in control neurons selectively phenocopies KCNQ2-DEE neuron dysfunction

We next sought to determine whether the functional phenotypes we identified in KCNQ2-DEE neurons reflected a specific consequence of impaired M-channel function rather than a general feature of epileptic or hyperexcitable iPSC-derived neurons. To test this, we chronically treated neurons from two mutation-corrected isogenic control lines (isoQ2-04^+/+^ and isoQ2-01^+/+^) with two different K^+^ channel blockers beginning on day 14 and assessed whether these treatments phenocopied the activity of corresponding KCNQ2-DEE neurons. We specifically treated neurons with the K_V_7 channel inhibitor XE991 (1μM) or with the non-specific K^+^ channel blocker 4-aminopyridine (4AP; 500μM), which does not inhibit K_V_7 channels, but has been shown to promote neuronal hyperexcitability and seizure-like activity in both animal models and iPSC-derived neurons^68-71^. We used our unbiased machine learning-based functional phenotyping platform to assess the impact of drug treatment, focusing on the three most predictive functional biomarkers of KCNQ2-DEE: spike count Fano factor, ISI CV, and percentage of spikes occurring within bursts (Figure 4E). We found that chronic XE991 treatment of mutation-corrected isogenic control neurons caused pronounced increases in all three KCNQ2-DEE biomarkers (Figure 6A). Notably, the XE991-induced phenotype was consistent across both patient genetic backgrounds and exceeded the effects observed in KCNQ2-DEE neurons, likely due to a more complete blockade of M-current compared to heterozygous loss-of-function *KCNQ2* mutations (Figures 6A and S10A). In contrast, general K^+^ channel blockade by 4AP did not phenocopy the features of patient neurons (Figure 6A). Instead, 4AP initially caused a transient increase in burst duration, number of spikes per burst, and median ISI within bursts but later, potentially through compensatory mechanisms, promoted greater network synchrony and elevated spike time tiling coefficient (Figure S10B).

**Figure 6.**
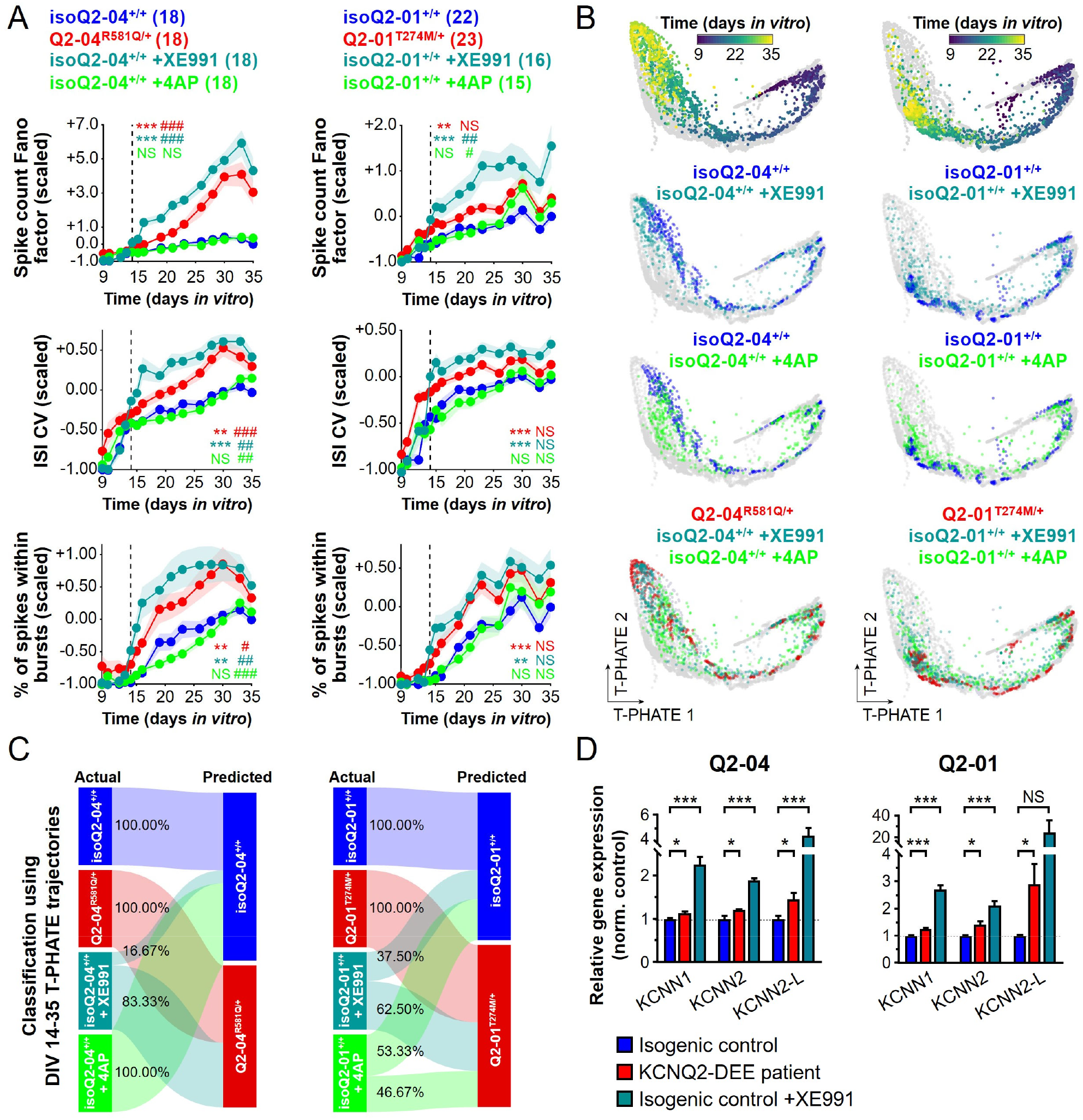
Machine learning predictions and gene expression analyses demonstrate that chronic K_V_7 inhibition selectively phenocopies KCNQ2-DEE functional and molecular signatures in isogenic control neurons. **A)** Chronic inhibition of K_V_7 channels with XE991 (1 µM applied from day 14) significantly increased the spike count Fano factor, ISI CV, and the percentage of spikes occurring within bursts, previously identified as the most predictive functional biomarkers of KCNQ2-DEE across time. These changes indicate greater firing irregularity and higher bursting propensity in XE991-treated isogenic control neurons. In contrast, chronic blockade of other K^+^ channels with 4-aminopyridine (4AP, 500 µM) did not modify the spike count Fano factor, ISI CV, or percentage of spikes within bursts in either isoQ2-04^+/+^ (left) or isoQ2-01^+/+^ (right) cultures. Repeated-measures ANOVA on drug treatment days 14-35: genotype effect: *p < 0.05, **p < 0.005, ***p < 0.0005; genotype/time interaction: # p < 0.05, ## p < 0.005, ### p < 0.0005; NS: not significant. Colored symbols indicate significant differences between isogenic control and KCNQ2-DEE neurons (red), XE991-treated isogenic control (teal), and 4AP-treated isogenic control (green) neurons. Data are shown as mean ± SEM, and the number of replicate wells is indicated in the panel. **B)** Functional embeddings of pharmacologically treated replicates of isoQ2-04^+/+^ (left) and isoQ2-01^+/+^ (right) neurons were projected onto the T-PHATE latent space to reveal altered functional trajectories. **C)** Supervised classifiers trained to distinguish the functional trajectories of KCNQ2-DEE neurons from those of their respective isogenic controls using T-PHATE embeddings from days 14-35 were used to assess the impact of XE991 and 4AP treatment on functional profiles of neurons. XE991 shifted the trajectory of isoQ2-04^+/+^ neurons toward a KCNQ2-DEE profile, whereas isoQ2-01^+/+^ neurons were less affected. While 4AP did not alter the trajectories of isoQ2-04^+/+^ neurons, it led to approximately half of the isoQ2-01^+/+^ neurons being classified as KCNQ2-DEE. **D)** Comparison of qPCR gene expression of SK channels among untreated isogenic control, KCNQ2-DEE patient, and XE991-treated isogenic control neurons at day 35. Expression of *KCNN1, KCNN2*, and *KCNN2-L* was significantly higher in KCNQ2-DEE patient neurons compared to respective isogenic controls (t-test; Q2-04: *p=0.0252, *p=0.0257, and *p=0.0281; Q2-01: ***p=0.0002, *p=0.0154, and *p=0.0306 for *KCNN1, KCNN2*, and *KCNN2-L* respectively). Expression of *KCNN1, KCNN2* and *KCNN2-L* was significantly higher in chronically XE991-treated isogenic control neurons compared respective untreated isogenic controls (t-test; Q2-04: ***p<0.0001, ***p<0.0001, and ***p=0.0003; Q2-01: ***p<0.0001, ***p<0.0001, and not significant p=0.0732 for *KCNN1, KCNN2*, and *KCNN2-L*, respectively). Data are shown as mean ± SEM from 3 independent differentiations with 2 technical replicates, normalized to 3 housekeeping genes and within each differentiation to respective isogenic controls. NS: not significant.

To determine the impact of XE991 and 4AP on the functional trajectory of iPSC-derived neurons, we projected pharmacologically-treated wells onto the lower-dimensional T-PHATE space (Figure 6B). Within each genetic background, we trained supervised classifiers on T-PHATE embeddings between days 14 and 35 to distinguish KCNQ2-DEE neurons from isogenic controls. We subsequently used these trained models to classify drug-treated neurons as either mutant or control (Figure 6C). This analysis showed that chronic K_V_7 channel inhibition with XE991 shifted the developmental trajectories among 83.3% of isoQ2-04^+/+^ and 62.5% of isoQ2-01^+/+^ mutation-corrected isogenic control neurons towards those of corresponding KCNQ2-DEE neurons (Figure 6C). In contrast, 4AP-treated neurons were classified as KCNQ2-DEE neurons in 0% of cases for isoQ2-04^+/+^, and 46.6% of cases for isoQ2-01^+/+^ (Figure 6C). These data-driven predictions suggest that M-current inhibition in isogenic control neurons reliably phenocopies functional features of KCNQ2-DEE neurons, whereas inhibition of other K^+^ channels does not. In agreement with this analysis, we found that XE991 treatment caused a dramatic increase in the expression of SK channels (*KCNN1, KCNN2, KCNN2-L*) in mutation-corrected isogenic control neurons (Figure 6D), demonstrating that blocking M-channels is sufficient to induce the dyshomeostatic upregulation of SK channels that we observed among neurons derived from all KCNQ2-DEE patient iPSC lines (Figure 1E-F). Collectively, these results demonstrate that early and chronic M-current inhibition is sufficient to recapitulate both the functional and molecular hallmarks of KCNQ2-DEE, reinforcing the specificity of our machine learning-based disease biomarkers.

### Retigabine rescues functional and molecular KCNQ2-DEE neuron phenotypes with variable efficacy

The utility of our machine learning analysis platform lies in its ability to predict patient-specific responses to therapeutic interventions. Retigabine is a positive allosteric activator of K_V_7 channels that has previously been used as an antiseizure medication to treat KCNQ2-associated epilepsy patients with variable efficacy^72-74^. Given that all 5 variants were diminish KCNQ2 current density or alter its activation voltage-dependence^10^, we treated patient iPSC-derived neurons with retigabine (5μM, days 14-35) and asked whether chronic K_V_7 activation would rescue KCNQ2-DEE functional phenotypes. We found that treatment with retigabine restored spike timing regularity and suppressed bursting propensity (as measured by spike count Fano factor, ISI CV, % of spikes within bursts) in neurons derived from all 5 KCNQ2-DEE patient iPSC lines (Figures 7A and S11A). Notably, retigabine also reduced the number of active neuronal populations in all 5 KCNQ2-DEE lines as well as the WMFR in 4 out of 5 lines (Figure S11B). To assess the impact of K_V_7 activation on the functional trajectory of KCNQ2-DEE neurons, we applied the T-PHATE-based classification analysis (Figure 7B). Chronic retigabine treatment shifted the trajectories of KCNQ2-DEE neurons towards those of their mutation-corrected isogenic controls with variable efficacy across lines, ranging from 100% in Q2-04^R581Q/+^ to 61.1% for Q2-17^H228R/+^ (Figure 7C). Accordingly, we found that while chronic retigabine treatment significantly reduced *KCNN1* and *KCNN2-L* expression in Q2-04^R581Q/+^ neurons, it only suppressed *KCNN2* expression in Q2-01^T274M/+^ neurons and had no effect on Q2-03^R207W/+^ and Q2-07^P335L/+^ neurons (Figure 7D). Importantly, in retigabine-treated Q2-17^H228R/+^ neurons, which exhibited the lowest proportion of phenotypic reversion, *KCNN2* expression was higher compared to untreated cells (Figure 7D). These findings indicate that the ability of retigabine to reverse maladaptive SK channel gene expression is limited and highly patient-specific, suggesting variable sensitivity to M-channel activation among different KCNQ2-DEE genotypes.

**Figure 7.**
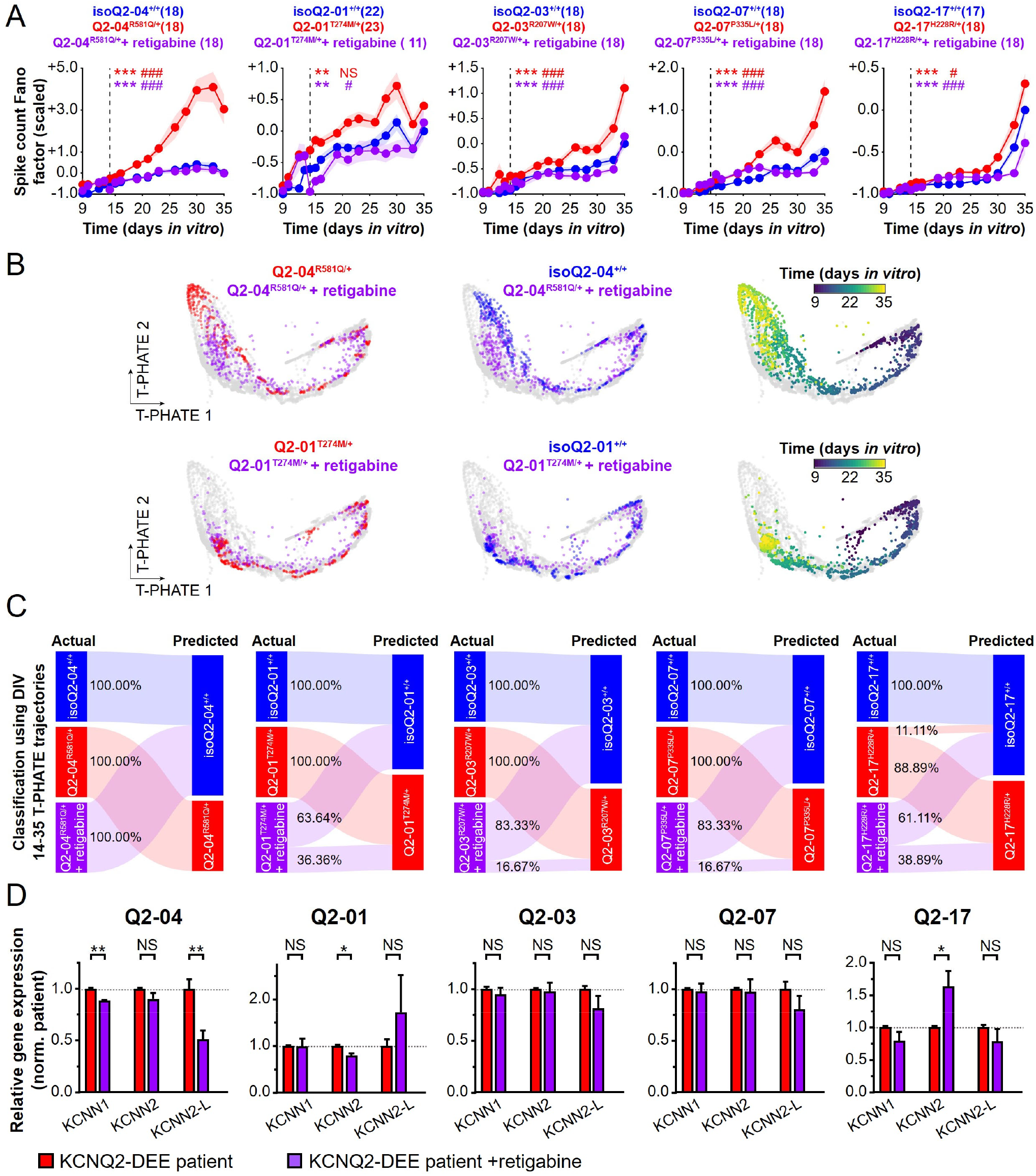
Machine learning predictions and gene expression analyses demonstrate that the K_V_7 channel activator retigabine rescues KCNQ2-DEE phenotypes with variable efficacy. **A)** Chronic activation of K_V_7 channels with retigabine (5 µM applied from day 14) reduced the spike count Fano factor, the most predictive functional biomarker of KCNQ2-DEE, promoting more regular firing patterns across all 5 KCNQ2-DEE patient iPSC neurons. Repeated-measures ANOVA on drug treatment days 14-35: genotype effect: *p < 0.05, **p < 0.005, ***p < 0.0005; genotype/time interaction: # p < 0.05, ## p < 0.005, ### p < 0.0005; NS: not significant. Colored symbols indicate significant differences between isogenic control and KCNQ2-DEE neurons, (red) and between KCNQ2-DEE and retigabine-treated KCNQ2-DEE neurons (purple). Data are shown as mean ± SEM, and the number of replicate wells is indicated in the panel. **B)** Functional embeddings of retigabine treated replicates of Q2-04^R581Q/+^ (top) and Q2-01^T274M/+^ (bottom) neurons were projected onto the T-PHATE latent space to evaluate changes in functional trajectories of KCNQ2-DEE neurons following retigabine treatment. **C)** Supervised classifiers previously trained to distinguish the functional trajectories of KCNQ2-DEE neurons from those of their respective isogenic controls using T-PHATE embeddings from days 14-35 were used to assess the impact of retigabine. Retigabine shifted the trajectories of Q2-04^R581Q/+^, Q2-03^R207W/+^, and Q2-07^P335L/+^ neurons toward control-like profiles, whereas Q2-01^T274M/+^ and Q2-17^H228R/+^ neurons exhibited weaker responses. **D)** Comparison of qPCR gene expression of SK channels among untreated and chronically retigabine-treated KCNQ2-DEE patient neurons at day 35. Chronic treatment with retigabine reduced expression of *KCNN1* and *KCNN2-L* in Q2-04^R581Q/+^ neurons (t-test; **p=0.0006 and **p=0.0045, respectively) and *KCNN2* in Q2-01^T274M/+^ (t-test; *p=0.0168) and increased expression of *KCNN2-L* in Q2-17^H228R/+^ neurons (t-test; *p=0.0326). Data are shown as mean ± SEM from 3 independent differentiations with 2 technical replicates normalized to 3 housekeeping genes and within each differentiation to respective KCNQ2-DEE neurons. NS: not significant.

## Discussion

KCNQ2-DEE is a devastating and currently untreatable neurodevelopmental disease characterized by early onset neonatal seizures and poor prognosis^1-3^. Our understanding of how pathogenic *KCNQ2* mutations impair the excitability of human neurons is limited. Using a diverse set of patient-specific iPSC-derived neurons for electrophysiological recordings and machine learning based data analysis, we identified a convergent pathophysiological signature of KCNQ2-DEE featuring irregular spike timing and enhanced bursting, as well as a disruption in the functional developmental trajectory of excitatory neurons. This dysregulation consistently correlated with transcriptional upregulation of SK channels accompanied by larger slow post-burst AHPs. Despite genetic heterogeneity, these shared electrophysiological and transcriptional phenotypes underscore a broad maladaptive compensatory mechanism transcending the primary K_V_7 channel dysfunction. In support of this model, we show that pharmacological inhibition of K_V_7 channels in mutation-corrected isogenic control neurons is sufficient to induce SK channel upregulation and the irregular firing patterns we found in the mutant KCNQ2-DEE neurons. Furthermore, although the K_V_7 activator retigabine rescued these disease-associated functional phenotypes and SK channel expression, it did so with variable efficacy across patients. Together, these findings establish maladaptive homeostatic regulation as a central determinant of KCNQ2-DEE pathogenesis with potential as a therapeutic target.

Increasing evidence points to dyshomeostatic compensation as a common pathophysiological theme across various neurodevelopmental epilepsy disorders^75,76^. In particular, upregulation of SK channel conductance has been implicated in learning and memory deficits^77-79^ and in several neurodevelopmental disorders. For instance, loss of *UBE3A* in Angelman syndrome elevates synaptic SK2 levels, impairing plasticity and cognition^79,80^. Similarly, *PTEN* haploinsufficiency promotes SK channel overexpression and enhanced post-burst AHPs, reducing cortical responsiveness^81^. Genetic mutations associated with autism and schizophrenia, notably truncation of *DISC1* and 16p11.2 deletion, also share synaptic phenotypes characterized by increased SK2 expression and prolonged spike refractory periods in mouse hippocampal and cortical neurons^82-84^. Additionally, modulation of SK channels has been shown to disrupt cytoskeletal organization and neuronal maturation^85^. These findings emphasize that altered SK channel conductance contributes to network dysfunction in diverse neurodevelopmental disorders, even when the primary mutation does not involve an ion channel. Our work extends this pattern to include KCNQ2-DEE, suggesting that maladaptive SK channel upregulation may represent a shared neuronal response to chronic perturbations in neuronal excitability. This convergence among distinct etiologies suggests that neurodevelopmental disorders may reflect disruptions in homeostatic mechanisms aimed at stabilizing neuronal activity, rather than exclusively resulting from the direct consequences of the initiating mutation.

Our results align with a growing body of work describing seemingly contradictory electrophysiological profiles in channelopathy-associated epilepsy. For example, loss-of-function mutations in *SCN1A* and *SCN2A* Na^+^ channels^86-89^ and gain-of-function mutations in K^+^ channels including K_V_1.2 (*KCNA2*), K_V_7.3 (*KCNQ3*), SK3 (*KCNN3*), BK (*KCNMA1*), and K_Na_1.1 (*KCNT1*)^90-95^, which are predicted to reduce neuronal excitability, have paradoxically been shown to cause hyperactivity in pyramidal neurons. One explanation for this apparent contradiction is that these effects reflect failures in autoregulatory feedback, where homeostatic rebalancing of ion channel expression^96^ and synaptic remodeling become maladaptive rather than stabilizing^75,76^. Our findings demonstrate that *KCNQ2* loss-of-function mutations lead to early onset of spontaneous activity and a cascade of maladaptive compensation that promotes aberrant neuronal bursting activity. Notably, burst-firing drives larger Ca^2+^ influx, which, due to the reliance of many cellular functions on Ca^2+^ homeostasis, profoundly affects neuronal maturation and both intrinsic and synaptic homeostatic plasticity^60,97-100^. This model may also help explain why KCNQ2-DEE patients experience both seizures and severe developmental delays.

The use of mutation-corrected isogenic control iPSC lines has become a gold standard in stem cell-based disease modeling studies^1^. To our knowledge, our work is the first to utilize 5 pairs of isogenic lines derived from patients with similar clinical presentations, yet distinct mutations in the same gene. Cellular models of genetic disorders that examine a single pathogenic mutation, leave uncertainty about whether observed phenotypes are variant-specific or reflect shared pathogenic pathways common to multiple mutations within the same disorder. Our findings reinforce and extend our previous observations with a single patient line^52^, establishing SK channel upregulation as a key pathophysiological mechanism and irregular spike timing as a functional biomarker of KCNQ2-DEE.

MEA technology offers a holistic, scalable assessment of the functional properties of populations of neurons, making it easier to interrogate neurons derived from multiple patient iPSC lines simultaneously^58,101^. However, relying on firing rate alone fails to capture the complexity of neural activity^102,103^. At the same time, analyzing the high-dimensional spatiotemporal data generated by MEA systems remains challenging, especially when experiments span many time points, recordings, and treatment conditions. This underscores the need for methods that can capture the diverse activity features of neuronal networks and reliably detect phenotypic differences over time *in vitro*.

The integrated computational framework we designed and implemented in this study addresses this critical gap. It allowed us to capture and interpret the complex functional signatures specific to KCNQ2-DEE cortical excitatory neurons, despite heterogeneous genetic backgrounds. Importantly, our platform leveraged T-PHATE dimensionality reduction to embed temporal and genotype-specific information, enabling robust visualization of disease trajectories over time. This analysis surprisingly revealed that the genetic background is a dominant factor in determining how neurons develop their firing patterns across time.

The machine learning analysis platform additionally allowed us to assess the effects of pharmacological treatments on neuronal function and predict patient-specific responses to therapeutic interventions. Activation of K_V_7 channels with retigabine produced inconsistent outcomes among KCNQ2-DEE patient iPSC neurons: some migrated toward their respective isogenic control state, whereas others remained considerably pathologic. Furthermore, retigabine led to reductions in SK channel transcript expression only in 2/5 cases. While these findings demonstrate the translational utility of our computational framework for guiding precision drug discovery in developmental epilepsy disorders, they also highlight the complexity of therapeutic responses even within a shared clinical diagnosis. Importantly, limited clinical data suggests that retigabine treatment of KCNQ2-DEE patients leads to variable responses, correlating earlier treatment with better outcomes^72-74^.

An important caveat of retigabine is its lack of specificity; in addition to activating K_V_7 channels, it functions as a GABA_A_ receptor agonist^104,105^, and inhibits K_V_2.1 channels^106^. Additionally, the overall reduction in firing rate in response to retigabine complicates the interpretation of spike count irregularity and bursting features, our most predictive KCNQ2-DEE biomarkers, because calculation of these features is highly sensitive to the underlying firing rate. The use of more selective K_V_7 channel activators, as well as initiating treatment earlier in neuronal development, may help clarify whether the variable efficacy of retigabine is due to delayed intervention, off-target effects, or the irreversibility of dyshomeostatic responses once triggered. Alternatively, bolstering M-current activation pharmacologically might not be a relevant approach for some *KCNQ2* mutations, which might be causing K_V_7 loss-of-function through altered stability or mislocalization of mutant channels. In such cases, alternative strategies such as allele-specific knockdown of mutant *KCNQ2* to alleviate dominant-negative effects or targeting dyshomeostaticly upregulated SK channels with anti-sense oligonucleotides (ASOs), might be rational approaches.

### Limitations of the study

While our study provides critical insights into KCNQ2-DEE disease mechanisms and patient-specific diversity, it also has important limitations. Our analysis focused exclusively on excitatory cortical neurons, representing a reductionist approach that isolates the impact of mutant K_V_7.2 channels on glutamatergic neurons, which are known to express high levels of these channels^107^. This strategy does not capture the broader effects of K_V_7.2 channel dysfunction within the full complexity of neural circuits. Future studies integrating inhibitory GABAergic neurons and additional neural cell types, potentially within the context of brain organoids or co-culture systems, may help promote greater understanding of how pathogenic *KCNQ2* mutations disrupt neuronal circuit dynamics.

Moreover, while our MEA-based computational and machine learning framework advances the current standard for functional analysis, our dataset is constrained by expert-curated feature selection. Future studies could leverage data-driven approaches, such as graph neural networks, to discover functional features directly from spike trains. Benchmarking the performance of alternative supervised machine learning algorithms, such as logistic regression and *k*-nearest neighbors, could help optimize classification performance and improve biomarker discovery. Importantly, our trained models are not static and can be continually refined with additional technical replicates from cell lines used in this study and from additional patient-specific lines harboring other pathogenic mutations in *KCNQ2* or other channels, thereby enhancing their translational robustness and utility.

Lastly, while our machine learning platform provided unbiased insights into therapeutic responses, the results should be interpreted with caution. Classifiers were constrained to assign pharmacologically treated culture wells to one of two predefined classes, either mutant *KCNQ2* or isogenic control neurons. Drug treatments may induce intermediate or entirely novel functional states that cannot easily be assigned to these categories. Future work could incorporate probabilistic classifiers to quantify prediction uncertainty and better reflect the spectrum of possible outcomes. Additionally, unsupervised learning approaches could uncover emergent functional clusters in treated neurons without imposing predefined labels, offering a more thorough understanding of drug-induced shifts in functional phenotypes.

## Acknowledgements

We are grateful to the following funding sources: US National Institutes of Health (NIH) National Institute on Neurological Disorders and Stroke (NINDS) R21NS125503 (D.S.), U54NS10887 (A.L.G. and E.K.), and the New York Stem Cell Foundation (E.K.). We would like to thank Shoai Hattori for providing and adjusting MATLAB protocols to analyze current-clamp data, Francesco Alessandrini for making the qPCR heatmaps and volcano plots, and Angel Alvarez at the Northwestern Stem Cell Core Facility for iPSC generation. E.K. is a New York Stem Cell Foundation – Robertson Investigator.

## Materials and Methods

### Clinical information on KCNQ2-DEE study participants

**Q2-04**^**R581Q/+**^: Subject is a female whose PBMC samples were collected at age 12 years. She has developmental delay and static encephalopathy with Lennox-Gastaut Syndrome secondary to pathogenic *KCNQ2* variant (R581Q). Seizure onset occurred shortly after birth and was described as body stiffening, jerking, with eyes deviated upwards, mouth pulled to one side, increased secretions and breath holding for 15 seconds to 1 minute. **Q2-01**^**T274M/+**^: Subject is a male whose PBMC samples were collected at age 7 years. He has epileptic encephalopathy with treatment-resistant epileptic spasms and myoclonic-tonic seizures secondary to a recurrent *KCNQ2* pathogenic variant (T274M). Seizures were first noticed on day 1 of life. Subject has developmental delay, plagiocephaly, congenital sternocleidomastoid torticollis, esotropia, congenital nystagmus, contracture of joint, optic atrophy, and bilateral chronic otitis media. Subject has gastrointestinal issues and constipation. Early myoclonic encephalopathy (EME) as an initial diagnosis, current diagnosis is clusters of epileptic spasms and brief tonic seizures.

**Q2-03**^**R207W/+**^: Subject is a female whose PBMC samples were collected at age 6 years. She has neonatal onset epilepsy with focal seizures secondary to a pathogenic *KCNQ2* variant (R207W). Seizures are partially controlled but persist despite anticonvulsant treatment. She has development delay, fasciculations of thigh muscles and myokymia. Neonatal MRI was unremarkable. Family history is significant for maternal neonatal onset epilepsy and same *KCNQ2* variant, brother with neonatal onset epilepsy now resolved.

**Q2-07**^**P335L/+**^: Subject is a male whose PBMC samples were collected at age 5 years. He had neonatal onset seizures with developmental delay secondary to pathogenic *KCNQ2* variant (P335L). Currently, he has features of autism without epilepsy in the absence of anti-seizure medications. **Q2-17**^**H228R/+**^: Subject is a female whose PBMC samples were collected at age 5 months. She had neonatal onset seizures secondary to pathogenic *KCNQ2* variant (H228R). There was burst-suppression documented by electroencephalogram (EEG). Seizures abated but subject has developmental delay.

### Generation, maintenance and CRISPR/Cas9 genome editing of human iPSC lines

Patient iPSC lines were generated from peripheral blood mononuclear cells (PBMCs) isolated from whole blood isolated from patients. Reprogramming of PBMCs into iPSCs was performed at the Northwestern Stem Cell Core Facility using Invitrogen’s CytoTune®-iPS 2.0 Sendai Reprogramming system (A16517, Thermofisher) as previously described^52,53^. All iPSCs were grown on Matrigel (BD Biosciences, BD354277) with mTeSR1 media (STEMCELL Technology, 85850), maintained at 37°C and 5% CO2 and passaged weekly using Accutase (Sigma). All cell lines were regularly tested for presence of mycoplasma using MycoAlert PLUS Detection Kit (Lonza) and determined to be mycoplasma-free.

Isogenic control iPSCs were generated using CRISPR/Cas9 from the respective patient-derived iPSC line in collaboration with Applied StemCell (Milpitas, CA). Briefly, one million patient iPSCs were electroporated with a mixture of guide RNA (gRNA) and Cas9 (in the ribonucleoprotein format), and ssODN. A small portion of the cell culture, presumably with mixed population, was subjected to Sanger sequencing analysis. Once the mixed culture showed repair with qualified HDR efficiency, the transfected cells were subjected to single cell cloning. Single cell-derived clones were cultured for 15 to 20 days, followed by genotype analysis by Sanger sequencing. Positive clones were further expanded and submitted again for sequencing to confirm desired genotype. iPSC clones were then cryopreserved and shipped to our lab. All gRNA and ssODN sequences as well as primer sequences used for PCR and Sanger sequencing for all of the patient and isogenic control lines have been previously published^52,53^.

For details on editing strategy, genomic DNA PCRs and Sanger sequencing, genomic integrity and pluripotency assays, analysis of off-target Cas9 sites and quantitative genotyping PCR-based copy number assays have been previously published^53^.

### Preparation of lentivirus

TetO-Ngn2-puro (Addgene plasmid #52047) and TetO-FUW-EGFP (Addgene plasmid #30130) plasmids were gifts from Marius Wernig^108,109^. FUW-M2rtTA (Addgene plasmid # 20342) was a gift from Rudolf Jaenisch^110^. Lentiviruses were generated in HEK293T cells using the second-generation packaging vectors, psPAX2 and pMD2.G, as described previously^111^ by the Northwestern University Gene Editing Transduction & Nanotechnology Core (GETiN).

### Generation of primary mouse glia

For all experimental analysis, iPSC-derived neurons were plated on primary mouse cortical glia, derived as previously described^112^. Primary glial cell cultures were derived from postnatal day 0-2, CD-1 mice (Charles River). Briefly, brain cortices were dissected free of meninges in dissection buffer HBSS (Thermo Fisher), then digested with trypsin (Thermo Fisher) and DNAse I (Worthington) for 10 min at 37°C. The tissue was dissociated in glia medium: DMEM (Corning, #15-013-CV) supplemented with Glutamax, D-glucose, 10% normal horse serum (Life Technologies), and penicillin-streptomycin (Thermo Fisher). After centrifugation and resuspension, cells were filtered through a 0.45-micron cell strainer and plated on poly-D-lysine coated plates with glia media at 37°C, 5% CO_2_ for 2 weeks. Afterwards, glial cultures were tested for mycoplasma, dissociated for expansion, and frozen in 10% DMSO/horse serum. All animal experiments were approved and conducted in accordance with the policies and guidelines set forth by the Northwestern University Institutional Animal Care and Use Committee (IACUC).

### Cortical excitatory neuron differentiation

iPSCs were differentiated into cortical glutamatergic neurons using a modified version of a protocol based on *Ngn2* overexpression^108,113^. Briefly, stem cells were dissociated as single cells using Accutase, re-suspended in mTeSR1 with 10 µM ROCK inhibitor (Y-27632, DNSK International, #129830-38-2), then incubated with lentiviruses (FUW-M2rtTA, TetO-Ngn2-Puro, TetO-FUW-EGFP) in suspension for 5 min before plating (100,000 cells/cm^2^). After 24 hours, lentivirus was removed and replaced with fresh mTeSR1 to expand transduced iPSCs. Cells were then passaged onto 10 cm plates for further expansion and subsequently replated at 100,000 cells/cm^2^ on multiple 10 cm plates. Differentiation was initiated the next day (Day 1) by switching to knockout serum replacement (KOSR) medium, composed of KnockOut DMEM supplemented with KSR, nonessential amino acids (NEAA), Glutamax (Life Technologies), 55 µM β-mercaptoethanol (Gibco; #21985023), 10 µM SB431542 (DNSK International), 100 nM LDN-193189 (DNSK International), 2 µM XAV939 (DNSK International), and 3 µg/ml doxycycline (Sigma). On Day 2, media was changed to a 1:1 mixture of KOSR and neural induction medium (NIM) containing DMEM:F12 supplemented with NEAA, Glutamax, N2 (Gibco, Life Technologies), 0.16% D-glucose (Sigma), 2 µg/ml heparin sulfate (Sigma), 3 µg/ml doxycycline, and 2 µg/ml puromycin (Sigma). On Day 3, cells were maintained in NIM with doxycycline (3 µg/ml) and puromycin (2 µg/ml). Neurons were cryopreserved on Day 4 in 10% DMSO/FBS (Hyclone FBS [VWR] or Gemini Stasis FBS [Gemini]).

For experimental analyses, cryopreserved neurons were plated onto primary mouse cortical glial monolayers. Glia were pre-plated on PDL/laminin-coated 6-well plates (qPCR) and coverslips (ICC and patch-clamp experiments), or PEI/laminin-coated MEA plates in glial medium consisting of MEM (Life Technologies) supplemented with Glutamax, 0.6% D-glucose, and 10% horse serum (Life Technologies). After 5-7 days, neurons were thawed (Day 5 post-induction) and plated directly onto glia in Neurobasal medium (NBM) supplemented with NEAA, Glutamax, N2, B27 (Life Technologies), BDNF (10 ng/mL; R&D Systems), 2% FBS (Hyclone or Gemini), 3 µg/ml doxycycline, and ROCK inhibitor. Half of the medium was replaced the next day (Day 6), and then every other day thereafter, using NBM supplemented with NEAA, Glutamax, N2, B27, BDNF (10 ng/mL), 2% FBS, and doxycycline (2 µg/ml).

### Immunocytochemistry

Neurons plated on coverslips were fixed with 3.7% formaldehyde (Sigma) in 4% sucrose/PBS for 15 minutes at room temperature, followed by three washes with cold PBS. Cells were then permeabilized and blocked simultaneously in PBS containing 0.1% Triton X-100 and 5% normal goat serum (NGS) for 1 hour at room temperature. Primary antibodies were applied overnight at 4°C in PBS with 5% NGS. The following primary antibodies were used: anti-GFP (Abcam ab13970, RRID: AB_300798; 1:10,000), anti-MAP2 (Millipore MAB3418, RRID: AB_94856; 1:1,000), and anti-vGLUT1 (Synaptic Systems 135 302, RRID: AB_887877; 1:200). The next day, coverslips were washed three times with cold PBS and incubated with secondary antibodies and DAPI (Invitrogen #33342; 1:1000) for 1 hour at room temperature. Secondary antibodies included Alexa 488 goat anti-chicken, Alexa 568 goat anti-mouse, and Alexa 647 goat anti-rabbit (Thermo Fisher Scientific; all 1:1000). All antibody dilutions were prepared in PBS containing 5% NGS. Following staining, cells were washed three times in PBS, briefly rinsed in distilled water, and mounted onto slides with ProLong Gold antifade reagent (Life Technologies). Neurons from the same differentiation batch were fixed and stained in parallel using identical antibody dilutions. Images for MAP2/GFP/vGLUT1-positive neuron quantification were acquired at 10× or 20× magnification using a Leica inverted Ti microscope.

### RNA isolation and qRT-PCR

Neurons were plated on a monolayer of mouse glial cells (500K neurons: 250K mouse glia per well). To inhibit post-mitotic proliferation, a short pulse treatment with 2 μM cytosine arabinoside (Ara-C) was administered in the feeding media on day 7 and removed on day 9, after which neurons were maintained with standard feeding protocols as previously described. Cells were harvested by scraping at the indicated time points following neuronal differentiation. RNA was extracted using the NucleoSpin RNA purification kit (Macherey-Nagel; #740955). Cells were lysed in RA1 buffer supplemented with β-mercaptoethanol, briefly vortexed, and cryopreserved according to the manufacturer’s protocol. First-strand cDNA was synthesized from 1.1 μg of DNase I-treated RNA (Invitrogen) using SuperScript™ IV Reverse Transcriptase (Thermo) and oligo(dT) primers following the manufacturer’s instructions. cDNA samples were diluted to correspond to 9.17 ng of input RNA per qPCR reaction. Quantitative PCR was performed using PowerUp SYBR Green Master Mix (Thermo; #A25742) on a CFX system (Bio-Rad). PCR cycling conditions were: 95°C for 3 minutes; 40 cycles of 95°C for 10 seconds and 60°C for 30 seconds; followed by a melt curve analysis from 65°C to 95°C, increasing by 0.5°C every 5 seconds. All reactions were run in duplicate. Gene expression was quantified using the ΔCt method: the average Ct of three housekeeping genes (GPI, CYC1, and RPLP0) was subtracted from the Ct of the target gene. Relative expression was calculated as 2^−ΔCt^ (ΔΔCt) and normalized to the indicated experimental control. All qPCR primers were human-specific, validated for efficiency, and a glia-only control was included for each primer set to confirm neuron-specific gene expression. The *KCNN2* gene encodes two SK2 splice isoforms. The longer isoform (*KCNN2-L*; NM_001372233.1), is under an alternative promoter, with a distinct N-terminal sequence that directs localization at glutamatergic postsynaptic densities where SK2 channels contribute to excitatory postsynaptic potentials (EPSPs)^114,115^. Our primers for *KCNN2* target both isoforms therefore, we included an additional set of primers targeting *KCNN2-L* specifically. All qPCR primer sequences are provided in Table S2.

### Patch clamp electrophysiology

Whole-cell current-clamp recordings were done as previously described^52^. We selected the earliest time point (days 22-26) where we previously detected enhanced post-burst AHPs in Q2-04^R581Q/+^ neurons and used current clamp electrophysiology to compare the intrinsic excitability properties of Q2-01^T274M/+^ and Q2-07^P335L/+^ patient neurons to their respective isogenic controls. Visually identified GFP-positive neurons were patched in whole-cell current-clamp mode under an Olympus IX51 inverted microscope fitted with a 40× objective. Glass micropipettes pulled on a Sutter P-1000 horizontal puller produced tip resistances of 2-4 MΩ when filled with a standard K-methyl sulfate internal solution (in mM): 120 K-MeSO_4_, 10 KCl, 10 HEPES, 10 Na_2_-phosphocreatine, 4 Mg-ATP, 0.4 Na_3_-GTP; pH 7.35 (KOH-adjusted), 285–290 mOsm/kg. Cells were continuously perfused (32-35 °C) with oxygenated aCSF containing (in mM): 125 NaCl, 26 NaHCO_3_, 2.5 KCl, 1.25 NaH_2_PO_4_, 1 MgSO_4_, 22 glucose, 2 CaCl_2_; pH 7.35, 310–315 mOsm/kg.

Signals were acquired with a Multiclamp 700B amplifier (Molecular Devices) and digitized at 10 kHz (3 kHz Bessel filter). Membrane potential was maintained at -65 mV (V_h_), and all reported voltages were corrected for an -8.2 mV liquid-junction potential. Resting membrane potential (RMP) was recorded immediately after break-in. Input resistance (R_N_) was derived from the slope of voltage responses to 500 ms current injections ranging from -50 pA to +30 pA in 10 pA increments. A 25-spike, 50 Hz train (2 ms, 1.2 nA suprathreshold pulses) elicited post-burst afterhyperpolarizations: the medium AHP (mAHP) was measured at the negative peak, and the slow AHP (sAHP) 1 second after train offset, both relative to holding potential. Single AP properties, including fast AHP (fAHP), were quantified using somatic current injection ramps (10-80 pA, 500 ms) and analysis of only the first AP elicited by each ramp. AP amplitude was defined as the difference between the AP peak and V_h_, whereas AP threshold was calculated where the first derivative of the up phase of the trace equaled 5 mV/ms. Using a 1 ms sliding window, fAHP was taken at the time dV/dt returned to 0 ± 0.5 mV/ms after initial spike in each sweep. AP half-width measurements were taken at half the AP peak amplitude relative to V_h_. Neurons meeting the following criteria were used: series resistance (R_S_) < 20 MΩ, membrane resistance (R_N_) > 200 MΩ, resting membrane potential (RMP) < -45 mV, and AP amplitude > 80 mV from V_h_. Data were analyzed using custom MATLAB protocols^52,116^. All MATLAB scripts are available for download at github.com/simkind/Patch-clamp-analysis.git (Simkin, 2021). Data collected at week 4 (days 22-26) were combined for statistical analysis using Statview software.

### Multi-electrode array (MEA) recordings

For MEA studies, 48-well CytoView MEA plates (Axion Biosystems) with 16 electrodes per well were coated with 0.1% PEI and laminin (20 μg/mL) according to manufacturer instructions. Primary mouse cortical glial cells were seeded at 40,000 cells/well and, 5-6 days later, 75,000 neurons/well (previously frozen on Day 4 of differentiation) were thawed and plated onto glial monolayer. Neurons were consistently plated on the same day of the week across experiments to align with a Monday/Wednesday/Friday feeding schedule, ensuring that recording days (days *in vitro*; DIVs) were consistent relative to media changes. Half of the media was exchanged on each feeding day, and spontaneous neuronal activity was recorded daily (except Sundays) using the Maestro 1 768-channel amplifier (Axion Biosystems) and AxIS v2.5 software. Recordings were conducted for 5 minutes at 37°C in 5% CO_2_/95% O_2_, followed by electrical stimulation consisting of 20 pulses at 0.25 Hz and 20 pulses at 0.5 Hz to promote neuronal maturation and migration toward the electrode field^117^. Signals were band-pass filtered (300-5000 Hz) with a Butterworth filter, digitized at 12.5 kHz/channel with a gain of 1200x, and spikes were detected using an adaptive threshold set to six times the root-mean-square (RMS) noise per electrode. Electrodes were considered active if they recorded ≥1 spike/min, and wells with fewer than 8 active electrodes on Day 35 were excluded from analysis. To capture spontaneous and media-induced firing dynamics, activity was analyzed every day from Day 9 to Day 16, and subsequently only on feeding days through Day 35.

For pharmacological studies, neurons were treated chronically with either XE991 (1 μM), retigabine (5 μM), or 4AP (0.5 μM) beginning on Day 14, with drugs included in all subsequent media changes through Day 35. Initial MEA experiments were conducted using 5 patient/isogenic control pairs, with three independent differentiations per pair, in which neurons were plated onto pre-established glial monolayers. These experiments generated the dataset used for functional characterization and biomarker identification (Figures 2-5). For the Q2-04 and Q2-01 lines, chronic drug treatments were performed in parallel with corresponding untreated wells during the original differentiations. For the Q2-03, Q2-07, and Q2-17 lines, additional MEA experiments (three independent differentiations per pair) were conducted in which neurons (45,000 cells/well) and glia (40,000 cells/well) were plated simultaneously to enhance neuronal concentration over the electrode field. In these experiments, cells were localized centrally using a 15 μL droplet, and 2 μM Ara-C was added from Day 7 to Day 9 to limit post-mitotic neuronal and glial proliferation. Although simultaneous plating modestly shifted the trajectory of neuronal maturation, the core functional differences between patient-derived and isogenic control neurons remained robust. To preserve consistency in modeling, only the original untreated dataset was included in the machine learning analysis to establish the functional phenotype of KCNQ2-DEE, while the additional experiments were used exclusively to assess pharmacological responses (Figure 6 and 7).

### MEA recording analysis and feature extraction

Spike analysis included calculation of weighted mean firing frequency (total spikes ÷ active electrodes ÷ 300 seconds) and inter-spike interval coefficient of variation (ISI CV; standard deviation ÷ mean ISI), which reflects short-timescale spike irregularity. In contrast, the spike count Fano factor, defined as the variance of spike counts across 6-second bins divided by the mean spike count^118^, was used to measure long-timescale variability in firing stability. Single-electrode bursts were detected using a Poisson surprise method with a surprise threshold minimum of 3. This algorithm is adaptive to the mean firing rate on each electrode according to a “surprise” threshold, adapted from^119^. The burst percentage (spikes within bursts ÷ total spikes × 100%), burst frequency (Hz; total bursts ÷ bursting electrodes ÷ 300 seconds), and inter-spike interval CV (ISI CV; SD ÷ mean IBI) were used to capture the temporal organization of neuronal firing, offering deeper insights into intrinsic neuronal development and excitability than mean firing rate alone.

Network bursts were detected using the Envelope method in the NeuralMetrics tool (Axion Biosystems). In brief, three network burst detection algorithms were available: (1) the ISI Threshold method, which identifies bursts based on fixed maximum inter-spike intervals and a minimum number of spikes across electrodes; (2) the Adaptive method, which dynamically adjusts the maximum inter-spike interval based on each well’s mean firing rate to account for continuous activity; and (3) the Envelope method, which detects bursts from peaks in a smoothed spike histogram exceeding a set threshold above baseline, thereby identifying discrete network burst boundaries more accurately. We selected the Envelope method because it more reliably delineates the start and end of network bursts compared to ISI Threshold and Adaptive methods, which often merge nearby spiking into artificially prolonged bursts, especially during immature stages when neurons fire sporadically. Although the Adaptive method performs well in detecting bursts during continuous firing by adjusting ISI thresholds dynamically, it can struggle to separate distinct bursts during low-activity periods. In contrast, the Envelope method better captures discrete bursting events as neurons transition from sporadic firing to coordinated network activity, which was critical for analyzing our cultures that had not yet reached full maturity. Ideally, combining Envelope and Adaptive detection would further improve accuracy. For Envelope detection, a threshold factor of 1.9, a minimum inter-burst interval of 100 ms, a minimum of 35% participating electrodes, and a burst inclusion threshold of 60% were applied. Additional average network burst parameters were extracted using a 0–2000 ms window with 1 ms bins to determine peak network burst firing rate, time to network burst peak firing rate, and spike count Fano factor within network bursts, providing measures of network synchrony and stability. In total, 36 functional features were extracted using the NeuralMetrics Tool (Axion Biosystems) and 5 additional features using custom MATLAB scripts. The additional features we extracted include mean ISI, spike time tiling coefficient^120^, spike count Fano factor within bursts, ISI CV within bursts, and mean inter-network burst interval. All features and their definitions are provided in Table S3. To capture spontaneous and media-induced firing dynamics, activity was analyzed every day from Day 9 to Day 16, and subsequently only on feeding days through Day 35.

### MEA computational pipeline

41 functional features were extracted for each culture well using the Neural Metrics Tool (Axion Biosystems) and custom MATLAB scripts (see Table S3 for a complete list of features and definitions), with any non-determined value set to 0. Culture wells with fewer than 8 active electrodes on the final day of recording (day 35) were excluded. Because neuronal feeding schedule introduced a confounding effect in MEA recordings, data were sampled from all recorded timepoints up till day 16, after which only recordings from feeding days were used. This approach yielded 15 total timepoints over a 27-day period. To reasonably mitigate sparsity in longitudinal recordings and ensure a complete time-series for each feature, values were linearly interpolated across timepoints within each culture well using the *interp1()* function in MATLAB. While interpolated data were used in downstream machine learning and analyses and visualization, they were excluded from statistical hypothesis testing.

To account for systematic batch effects introduced by independent iPSC differentiations, we performed within-batch feature scaling. Specifically, for each patient-isogenic control pair, the relative difference (*ŷ*) between each feature value (*y*) of each culture well (*n*) and a reference scalar constant (*c*) was computed at every timepoint as:

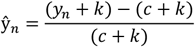

where *k* = 1 is included to prevent division by 0. The reference constant was defined as the mean feature value of isogenic controls on day 35. This effectively set feature means of isogenic controls to 0 on day 35, thus allowing comparability across features, differentiations, and patient-isogenic control pairs. To quantify time-resolved genotype effect sizes, we then subtracted the mean scaled feature value of isogenic controls from the corresponding scaled value of each culture well at each timepoint. This scaled differencing set the feature means of isogenic controls to 0 at all timepoints, enabling direct evaluation of feature up-or down-regulation in patient-derived neurons. Importantly, all feature scaling and scaled differencing steps were performed independently for each differentiation to minimize batch effects.

### Functional biomarker discovery

Disease-associated functional biomarkers of KCNQ2-DEE were identified using supervised machine learning. Supervised classifiers were trained to distinguish pooled patient-derived neurons from pooled isogenic controls at each timepoint, and feature importance scores from these models were used to identify functional biomarkers of KCNQ2-DEE across development. To ensure balanced representation of each cell line during pooling, scaled difference scores from 18 culture wells per cell line were sampled, which was the minimum sample size across all cell lines. To prioritize representative samples, we calculated how closely each culture well matched the average functional profile of its respective cell line. Each well was represented as a matrix ***X***:

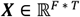

where, *F* = 41 is number of features and T = 27 is the number of timepoints. To quantify how representative each well was, we computed the root mean square deviation (RMSD) between its scaled difference feature matrix *X* and the cell line mean matrix 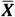 as follows:

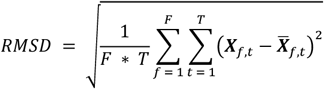

Culture wells with the lowest RMSD values were selected, ensuring that sampled wells best reflected the average functional profile of each cell line. Notably, since culture wells were sampled based on RMSD scores, independent differentiations may have contributed unequally to this down-sampled dataset. Sampled wells were then randomly assigned to training (16 wells per cell line; *n* = 160) or test (2 wells per cell line; *n* = 20) sets.

To identify functional biomarkers, ensemble classifiers composed of bagged decision trees were trained at each timepoint using the *fitcensemble()* function in MATLAB. The training set was split during 4-fold nested cross-validation to optimize performance and minimize overfitting. In each of 1,000 repetitions, the inner loop performed random search hyperparameter optimization by randomly sampling combinations from pre-defined ranges, while the outer loop assessed model generalization. Classifiers were trained on three (60 wells per class; *n* = 120) of the four folds using the optimized hyperparameters and validated on the remaining fold (20 wells per class; *n* = 40). This procedure was repeated across all folds, ensuring that every culture well in the training set contributed to both model training and evaluation. The table below lists the hyperparameters selected or optimized.

**Table.** Hyperparameters for ensemble classifiers.

| Parameter | Variable name | Optimization range or value |
| --- | --- | --- |
| Number of decision trees | *NumLearningCycles* | 20 to 200 |
| Maximum number of decision splits | *MaxNumSplits* | 0 to 2 |
| Minimum leaf size | *MinLeafSize* | 0 to 2 |
| Number of surrogate splits | *Surrogate* | 3 to 6 |
| Number of predictors to sample | *NumPredictorsToSample* | 41 |
| Predictor selection method | *PredictorSelection* | 'allsplits' |
| Split criterion | *SplitCriterion* | 'gdi' |
| Pruning method | *Prune / PruneCriterion* | 'on' / 'impurity' |
| Prior probability | *Prior* | 'uniform' |

Classifier performance during cross-validation was evaluated using the F_1_- score, which captures both sensitivity and specificity. F1-scores were computed as:

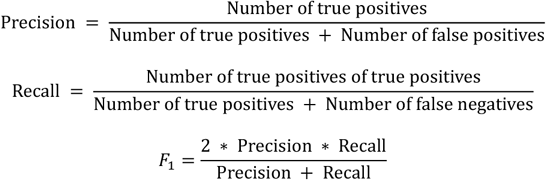

The combination of hyperparameters that yielded the highest median F_1_-score during the 4-fold nested cross-validation was selected to train a final model on the entire training set (*n* = 160). This optimized model was then evaluated on the held-out test set (*n* = 20) to assess its generalization performance.

To assess the contribution of individual features to classifier performance, a permutation-based feature importance analysis was performed using the *predictorImportance()* function. In this approach, the values of each feature were randomly shuffled across all culture wells in the training set, disrupting their association with the true class labels. A drop in classification performance following permutation indicated that the classifier relied on that feature for accurate predictions. To enable comparisons across timepoints, feature importance scores were normalized to the corresponding cross-validation F_1_-score, such that higher-performing classifiers yielded proportionally greater relative importance for informative features.

### Functional trajectory analysis

Functional trajectories of patient-derived and isogenic control neurons were uncovered using unsupervised machine learning. To ensure balanced representation of each cell line, scaled difference scores from 18 culture wells per cell line were sampled, as described above. The resulting high-dimensional array (*N* x 41), where *N* is the total number of culture wells across all timepoints (180 culture wells x 27 timepoints), was transformed using z-score standardization and reduced to a low-dimensional latent space (*N* x 3) using temporal potential of heat diffusion for affinity-based transition embedding (T-PHATE), implemented in Python. We used T-PHATE as other standard dimension reduction techniques including principal component analysis (PCA), *t*-distributed stochastic neighbor embedding (t-SNE), and unified manifold approximation and projection (UMPA), failed to capture the temporal dynamics in our dataset (Figure S9A). Each of these methods have limitations in the context of time-series data. PCA is a linear technique that cannot represent non-linear developmental trajectories; *t*-SNE focuses on preserving local structure within data at the expense of global relationships and temporal continuity; and, although UMAP better preserves global structure, it can disrupt smooth transitions across timepoints due to its reliance on nearest-neighbor graphs.

The T-PHATE (version 0.1) algorithm, specifically designed to capture temporal structure in time-series data, is publicly available at https://github.com/KrishnaswamyLab/TPHATE. T-PHATE was selected after initially testing more conventional dimensionality reduction techniques, including principal component analyses (PCA; implemented using the *pca()* function in MATLAB), *t*-distributed stochastic neighbor embedding (*t*-SNE; implemented using the *tsne()* function in MATLAB), and uniform manifold approximation and projection (UMAP; implemented using the *run_umap()* function in MATLAB with *min_dist* = 0.1, *n_neighbors* = 70, and *spread* = 1).

The table below lists the T-PHATE parameters, which were selected automatically.

**Table.**
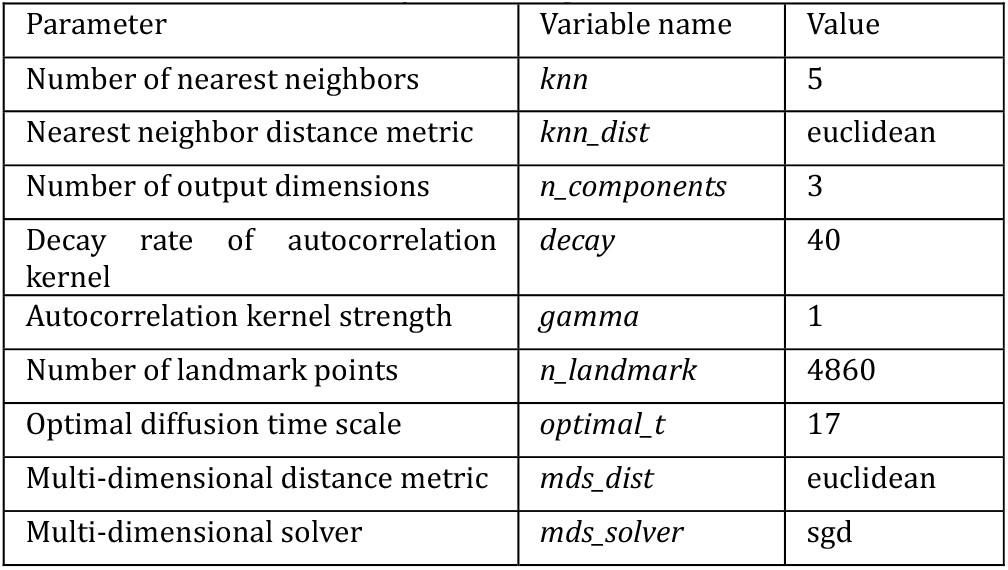
T-PHATE dimensionality reduction parameters.

To quantify spatiotemporal relationships between cell lines in the low-dimensional T-PHATE space, cluster analysis was performed using the Seurat package (version 5.1.0) in R. Specifically, the *FindNeighbors()* function was used to construct a *k*-nearest neighbor graph from the T-PHATE embeddings with a neighborhood size of 29. This graph was then transformed into a shared nearest neighbor graph, and weak edges (weight < 0.001) were pruned. Clusters were subsequently identified using the Louvain algorithm via the *FindClusters()* function with a resolution parameter of 1.

To assess whether cell lines within each genetic background exhibited distinct developmental trajectories, supervised machine learning classifiers were trained using the Statistics and Machine Learning Toolbox (version 24.2) in MATLAB. Specifically, the *fitcsvm()* function was used to train support vector machine (SVM) classifiers to distinguish patient-derived neurons from isogenic controls based on their day 9-35 T-PHATE embeddings. All hyperparameters, including box constraint, kernel function, and kernel scale, were optimized using Bayesian optimization, and classifier performance was evaluated using *k*-fold cross-validation with F_1_-score as the evaluation metric.

To assess the effects of small molecule drugs on developmental trajectories of neurons, supervised machine learning classifiers were first trained to distinguish untreated patient-derived neurons from untreated isogenic controls using their day 14-35 T-PHATE embeddings. Hyperparameters were optimized using Bayesian optimization, and classifier performance was evaluated using *k*-fold cross-validation with F_1_- score as the evaluation metric. To quantify drug-induced changes in developmental trajectories, treated culture wells were z-score standardized using parameters (mean and standard deviation) from the original *N* x 41 high-dimensional dataset and subsequently projected onto the previously computed low-dimensional T-PHATE space. Classifications were performed on the day 14-35 T-PHATE embeddings of treated wells. Importantly, culture wells in both training and test sets included neurons differentiated and plated using the same *Ngn2* excitatory cortical neuron differentiation and plating protocols, respectively.

### Spike sorting

Spike sorting is the computational process of grouping extracellular spike waveforms into distinct clusters, where each cluster is traditionally assumed to represent the activity of a single neuron. In this study, spike sorting was used as a quality control step to estimate the number of putative neurons per electrode per culture well on the final day of recordings (day 35).

Spike sorting was performed using the Statistics and Machine Learning Toolbox (version 24.2) in MATLAB. For each electrode, spike waveforms were transformed using z-score standardization and subsequently reduced to 3 principal components using the *pca()* function. Electrodes with fewer than 5 spikes were excluded from further analysis. The resulting principal component scores were clustered using *k*-means clustering with *k* ranging from 1 to 5. The optimal number of clusters was determined using the *evalclusters()* function based on the gap statistic and silhouette criterion. Single-cluster solutions were retained only if the gap statistic exceeded 0.75. For multi-cluster solutions, the configuration with the highest average silhouette value was selected. Importantly, all clustering solutions were manually inspected prior to final confirmation.

### Drugs

Drugs were prepared as stock solutions using distilled water or DMSO and then diluted to the required concentration in culture media immediately before use.

### Statistical analysis

For ICC, qPCR, patch-clamp studies, differences were evaluated using t-test, one-way, or two-way ANOVA where appropriate. For MEA data, repeated measures ANOVA was used to assess time-varying differences in functional

features; when assumptions of sphericity were violated, *p*-values were adjusted using lower-bound corrections. To control for multiple comparisons across features, *p*-value thresholds for statistical significance were adjusted using Bonferroni correction. All data are reported as mean ± SEM.

### KCNQ2-DEE patient study approval

Written informed consent was received from participants prior to inclusion in the study under protocols approved both by Ann & Robert H. Lurie Children’s Hospital of Chicago and Northwestern University IRB (#2015-738).

### Data availability

Extracellular spike voltages and spike times recorded using the Axion Maestro system (Axion Biosystems) and all original code for MEA recording analysis will be made publicly available at the time of publication.

## SUPPLEMENTARY TABLES AND FIGURES

**Figure S1.**
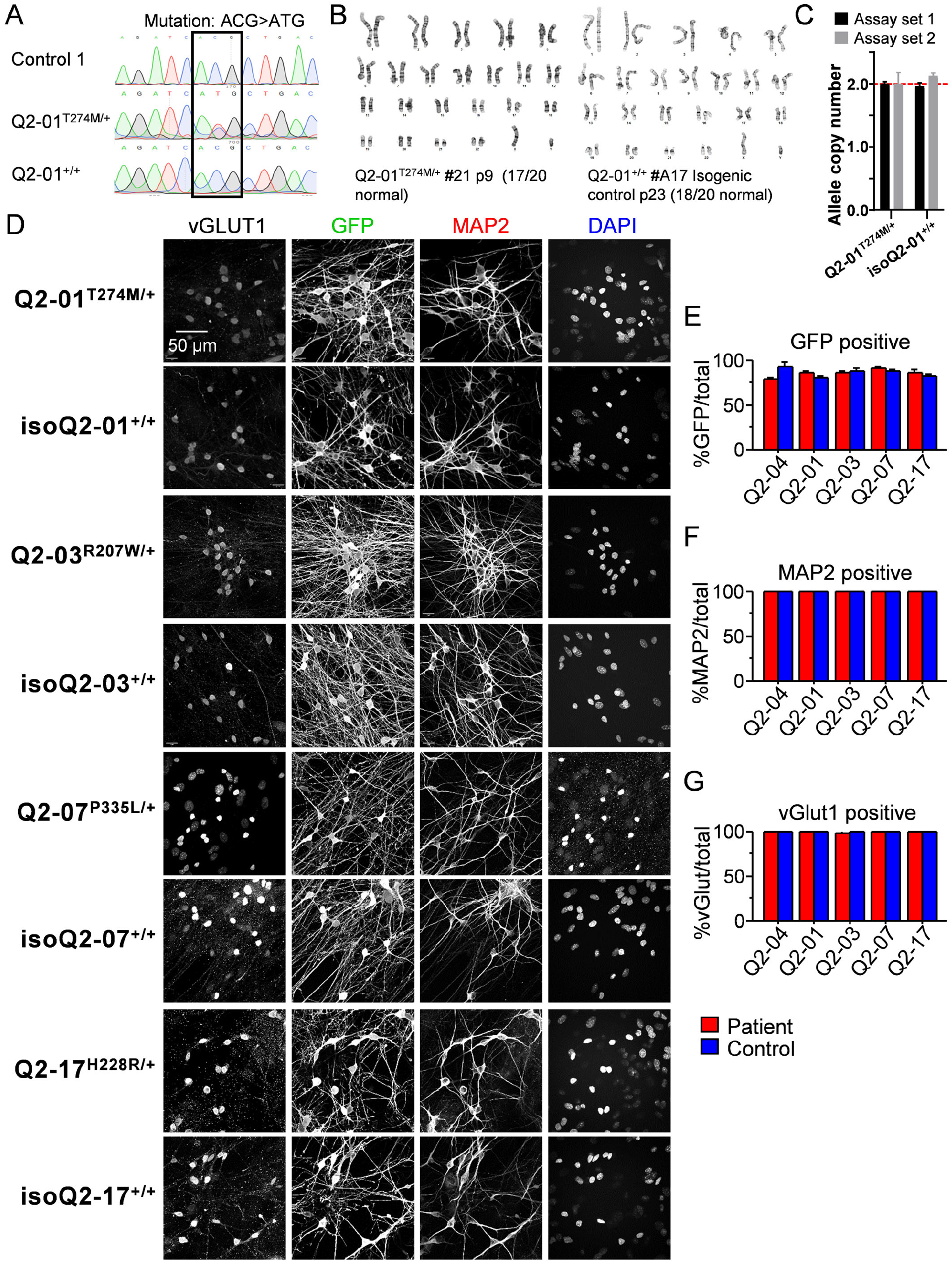
Validation of iPSCs and efficiency of cortical excitatory neuron differentiation. **A)** PCR amplification of on-target *KCNQ2* region for Q2-01^T274M/+^ patient line and Sanger sequencing electropherograms in control and patient iPSCs before and after gene editing, demonstrate the correction of the heterozygous (p.T274M; c.821C>T) in isogenic control isoQ2-01^+/+^ iPSCs. **B)** G-band karyotype analysis of Q2-01^T274M/+^ patient and CRISPR/Cas9 edited mutation-corrected isogenic control isoQ2-01^+/+^ lines revealed normal karyotypes. **C)** Genomic quantitative PCR (gqPCR) was used to assess allele copy numbers in Q2-01 isogenic iPSC lines, revealing bi-allelic expression using 2 different primer/probe assay sets. **D)** Immunocytochemistry of neurons derived from patient-specific KCNQ2-DEE iPSC lines and matched isogenic controls at day 26, labeled with neuronal (MAP2), glutamatergic (vGLUT1), and transgene expression (GFP) markers. DAPI staining marks cell nuclei. Scale bar: 50 µm. **E)** Quantification of GFP-positive neurons as a percentage of total identified neurons (calculated based on either GFP, MAP2, or vGLUT1 positive labeling) shows consistent differentiation efficiency across all KCNQ2-DEE and isogenic control pairs. Although not all neurons expressed detectable GFP (≥79%), no differences within pairs were observed. **F)** ≥99% of all neurons expressed MAP2, confirming robust neuronal differentiation with no significant differences between paired lines. **G)** ≥98% of all neurons expressed vGLUT1, indicating efficient differentiation into glutamatergic neurons, again with no significant differences between patients and their matched isogenic control lines. Data are shown as mean ± SEM.

**Figure S2.**
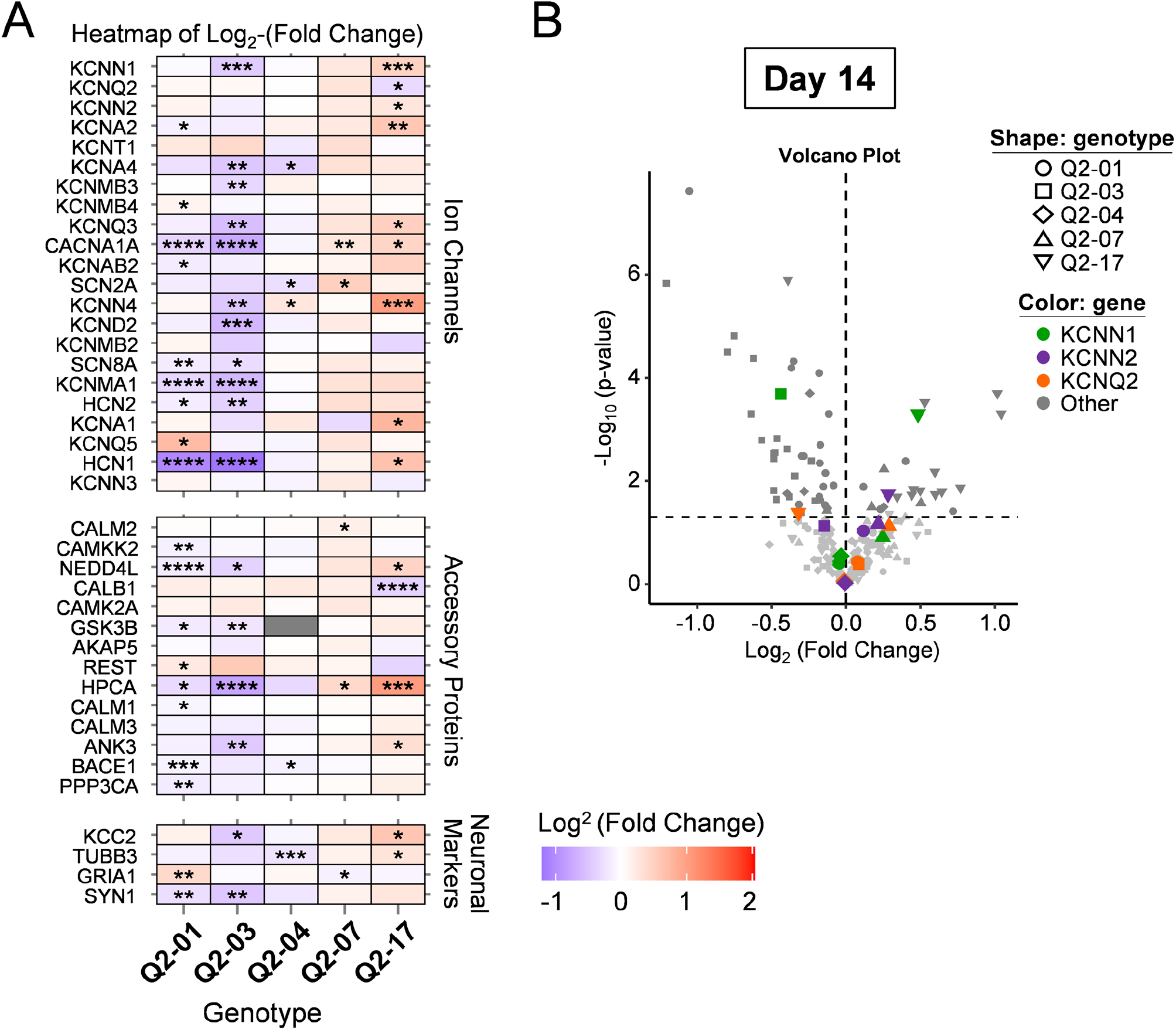
Gene expression profiles of KCNQ2-DEE neurons relative to respective isogenic controls at day 14. **A)** Heatmap of day 14 neuron gene expression profiles across 5 KCNQ2-DEE patient iPSC lines shown as log_2_ fold-change relative to their respective isogenic controls; N = 6 (3 independent differentiations x 2 technical replicates). Forty-two gene targets are first grouped by class (ion channels, accessory proteins, neuronal markers) and ranked within each class by the average log_2_ (p-values) x sign of fold change, showing significant fold-change up-regulation (top) or down-regulation (bottom). Red = up-regulated, purple = down-regulated; grey = not tested. Housekeeping genes: GPI, CYC1, RPLPO. t-test: *p < 0.05, **p < 0.005, ***p < 0.0005. **B)** Volcano plot displaying differential gene expression at day 14, with -log10(p-value) plotted against log2(fold-change). Different KCNQ2-DEE lines are denoted with different shapes. KCNN1, KCNN2, and KCNQ2 are highlighted in distinct colors. Horizontal dashed line marks the 5% significance threshold; vertical dashed line indicates ±1-fold (log_2_ = ±1).

**Figure S3.**
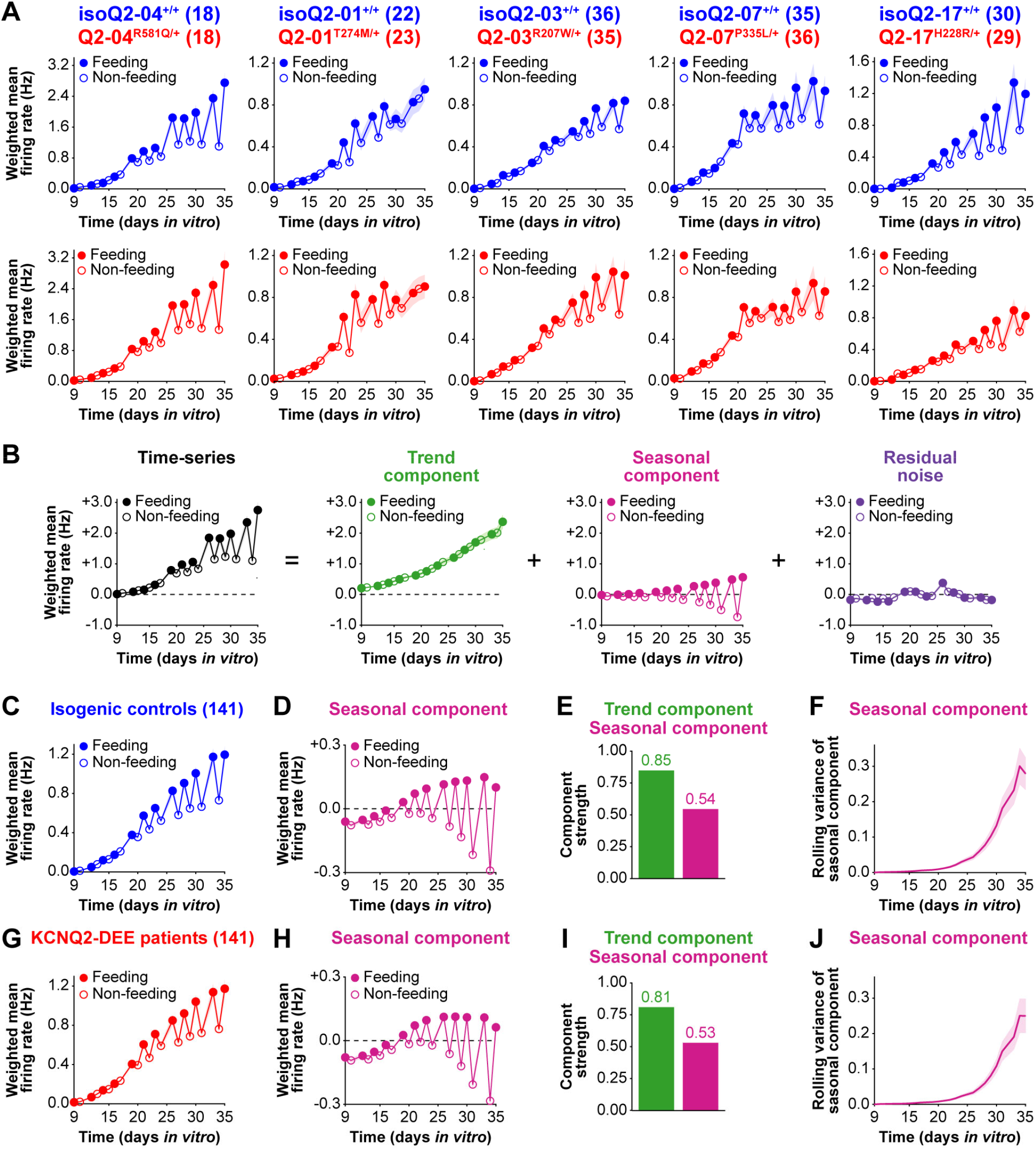
Neuronal feeding schedule introduces confounding effects in MEA recordings of iPSC-derived neurons. **A)** Daily weighted mean firing rates (WMFRs) of individual isogenic control (top) and KCNQ2-DEE (bottom) neurons recorded from days 9-35 show periodic fluctuations that align with the neuronal feeding schedule. Filled circles denote feeding days (recordings made ∼6 hours after media change), and open circles denote non-feeding days (recordings made ∼30 hours after media change). A pronounced post-feeding spike in activity is evident after day 16. **B)** Schematic illustrating time-series decomposition of WMFR using MATLAB’s trenddecomp() function, which applies singular spectrum analysis (SSA) to separate trend, seasonal, and residual components of signals. **C)** and **G)** Longitudinal analysis of WMFR across all isogenic control **(C)** and KCNQ2-DEE **(G)** neuronal culture wells. **D)** and **H)** Seasonal components extracted from isogenic control **(D)** and KCNQ2-DEE **(H)** neurons show cyclical shallow peaks on feeding days and deeper troughs on non-feeding days beginning around DIV 19, suggesting that non-feeding has a more dramatic suppressive effect on neuronal firing than the additive effect from feeding. **E)** and **I)** Trend and seasonal components show strong (0.81-0.85) and moderate (0.53-0.54) effect sizes, respectively in controls **(E)** and KCNQ2-DEE neurons **(I)**, indicating that feeding introduces a systematic confounding effect of top of the effect of neuronal maturation. **F)** and **J)** Rolling variance of the seasonal component exhibits sigmoidal growth over time in both isogenic control **(F)** and KCNQ2-DEE neurons **(J)**. To minimize confounding effect of feeding schedule, all recordings through day 16 were retained, after which only recordings made on feeding days were used for downstream analyses (see Methods). Data are shown as mean ± SEM.

**Figure S4.**
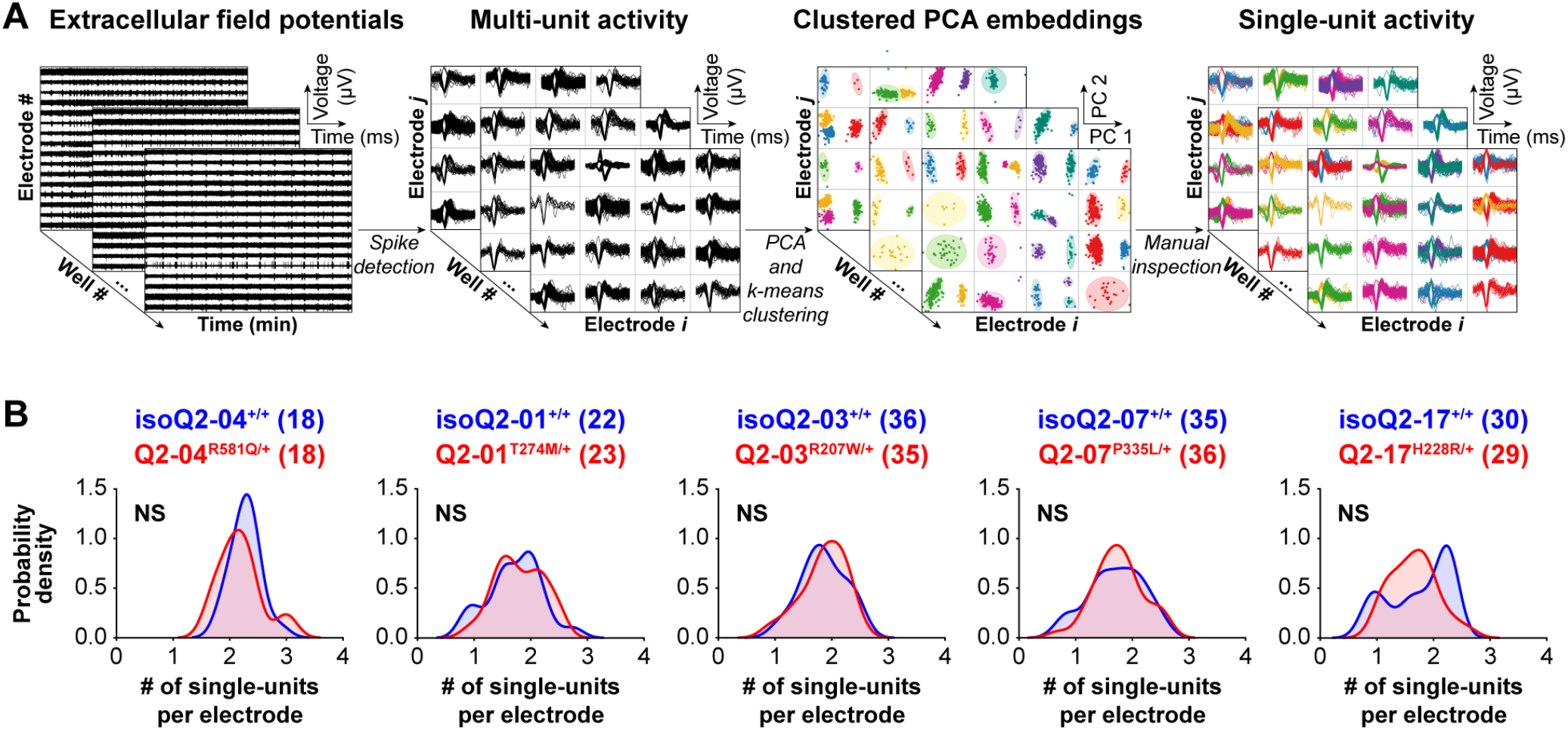
Unsupervised spike sorting demonstrates comparable neuron counts per electrode in KCNQ2-DEE and isogenic control cultures. **A)** Schematic of the spike sorting workflow used to identify single-units in 5 minutes of extracellular recordings on day 35. Spontaneous extracellular field potentials recorded from each micro-electrode of each well were thresholded to detect multi-unit spike waveforms. Multi-unit spike waveforms recorded on each microelectrode were then reduced to three principal components, which were clustered using k-means clustering to isolate putative single-units (right). **B)** Smoothed probability density plots show the distribution of the number of single-units per active electrode on day 35 for each patient-isogenic control pair. Across 8,236 single units (∼1.83 ± 0.03 single units per electrode), there was no statistically significant difference in the distribution of single-units per microelectrode between KCNQ2-DEE neurons and isogenic controls. These findings suggest that the early increase in active electrodes observed in KCNQ2-DEE cultures (Figure 2C) is unlikely to arise from different neuron density on the MEA but rather reflects intrinsically accelerated functional maturation. NS: not significant.

**Figure S5.**
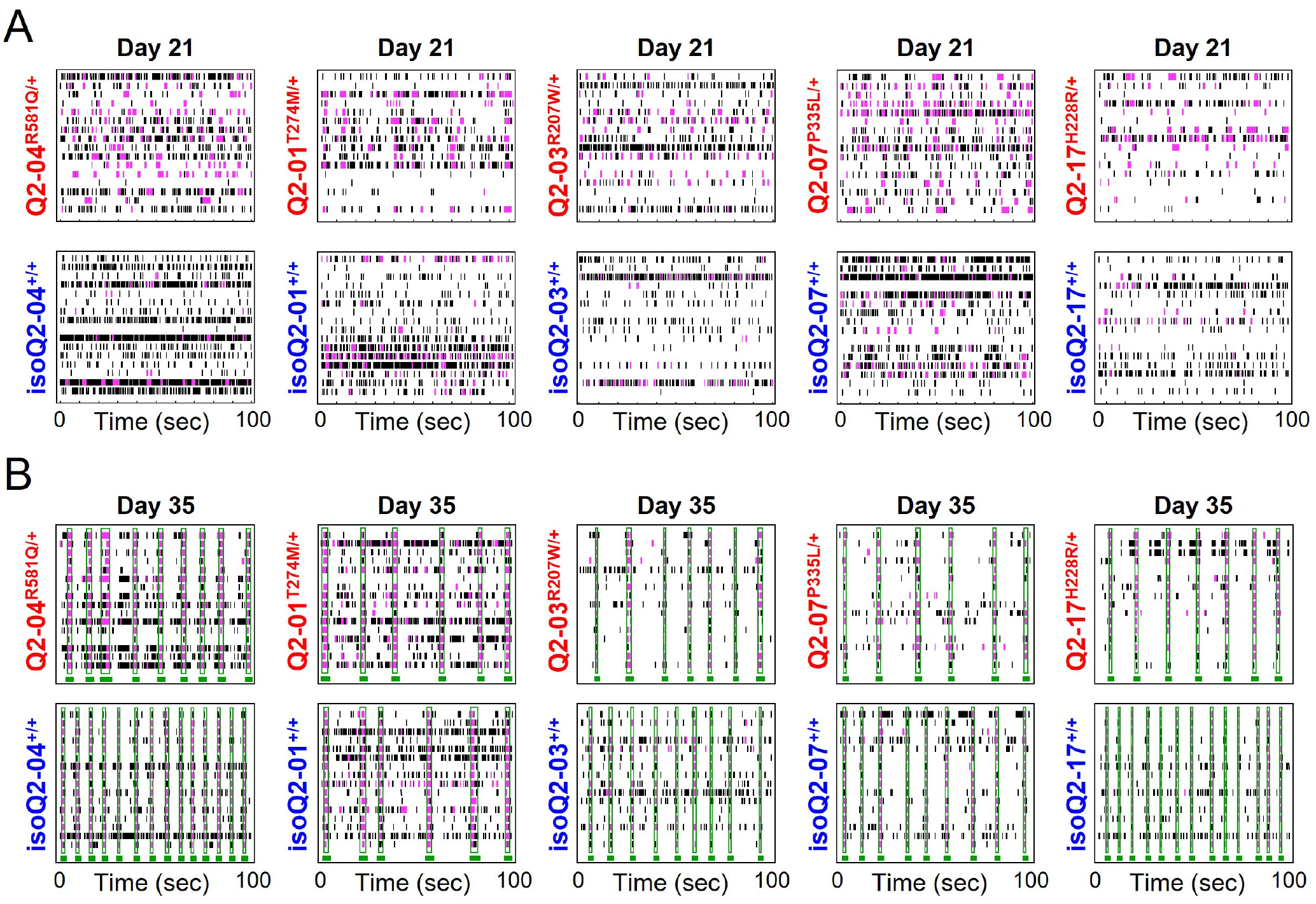
MEA activity raster plots reveal progression from single-electrode to network-level bursting in KCNQ2-DEE neurons. **A)** Representative 100-second raster plots of spiking activity on day 21 for each patient (top) and matched isogenic control (bottom). Each row represents one of 16 electrodes per well; black ticks indicate single spikes, and pink ticks denote spikes identified within single-electrode bursts. KCNQ2-DEE neurons show denser pink segments, reflecting the elevated burst percentage and spike-time irregularity quantified in Figure 2E-F. **B)** Representative 100-second raster plots of spiking activity raster plots on day 35 for each patient (top) and matched isogenic control (bottom). Network-level bursts, which are coordinated bursting across multiple electrodes are outlined in green.

**Figure S6.**
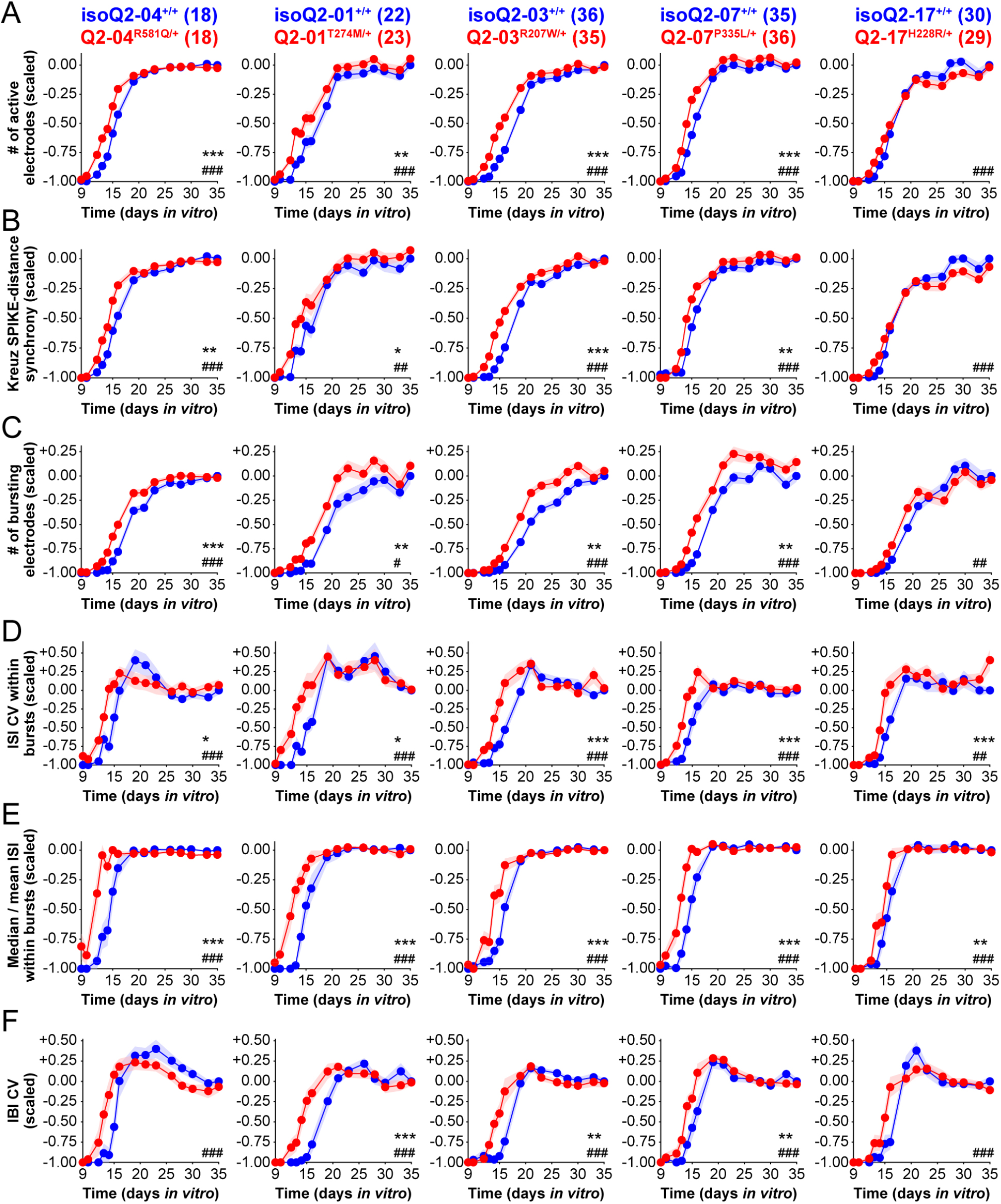
Early developmental divergence in firing properties across patient-specific KCNQ2-DEE iPSC neurons. Longitudinal analysis of six firing features that were consistently altered across all 5 KCNQ2-DEE lines at early developmental timepoints (days 9-20) compared to their respective isogenic controls. **A)** KCNQ2-DEE neurons recruited a greater number of active electrodes between days 9 and 20, indicating an earlier onset of active neuronal populations. **B)** KCNQ2-DEE neurons exhibited higher Kreuz SPIKE-distance synchrony values in the early time window, reflecting earlier emergence of synchronous activity within active neuronal populations. **C)** A greater number of electrodes exhibited bursting in KCNQ2-DEE neurons, suggesting an increased fraction of bursting neurons. **D)** The ISI CV within bursts was higher in KCNQ2-DEE neurons, indicating less regular spike timing during bursts. **E)** KCNQ2-DEE neurons showed higher median/mean ISI within bursts, suggesting an impaired ability to slow down during bursting in early developmental stages. **F)** Inter-burst interval (IBI) CV was elevated in patient-derived neurons, reflecting greater variability in timing between successive bursts. Repeated-measures ANOVA: genotype effect: *p < 0.05, **p < 0.005, ***p < 0.0005; genotype/time interaction: # p < 0.05, ## p < 0.005, ### p < 0.0005. Data are shown as mean ± SEM, and the number of replicate wells is indicated in the panel.

**Figure S7.**
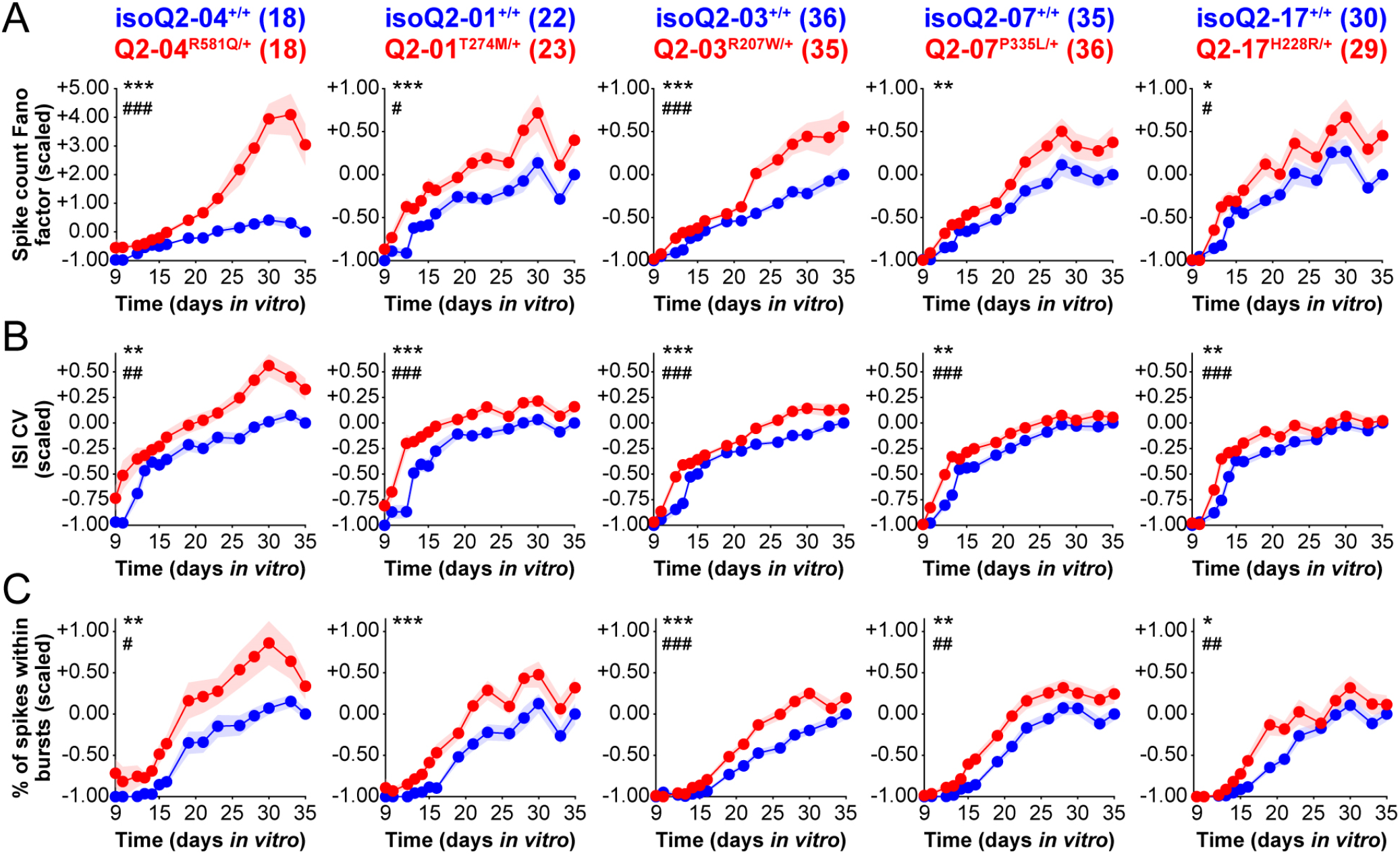
Longitudinal analysis of three firing pattern regularity features consistently altered across all 5 patient-specific KCNQ2-DEE iPSC neurons throughout the recording time course. **A)** KCNQ2-DEE neurons exhibit a higher spike count Fano factor across the recording period compared to their respective isogenic controls, indicating reduced firing regularity across time bins. **B)** ISI CV is elevated in KCNQ2-DEE neurons, reflecting greater variability in the timing between consecutive spikes and less precise spike timing. **C)** KCNQ2-DEE neurons display a higher percentage of spikes occurring within bursts, suggesting an overall increase in bursting propensity. Repeated-measures ANOVA: *p < 0.05, **p < 0.005, ***p < 0.0005 genotype effect. # p < 0.05, ## p < 0.005, ### p < 0.0005 genotype/time interaction. Data are shown as mean ± SEM, and the number of replicate wells is indicated in the panel.

**Figure S8.**
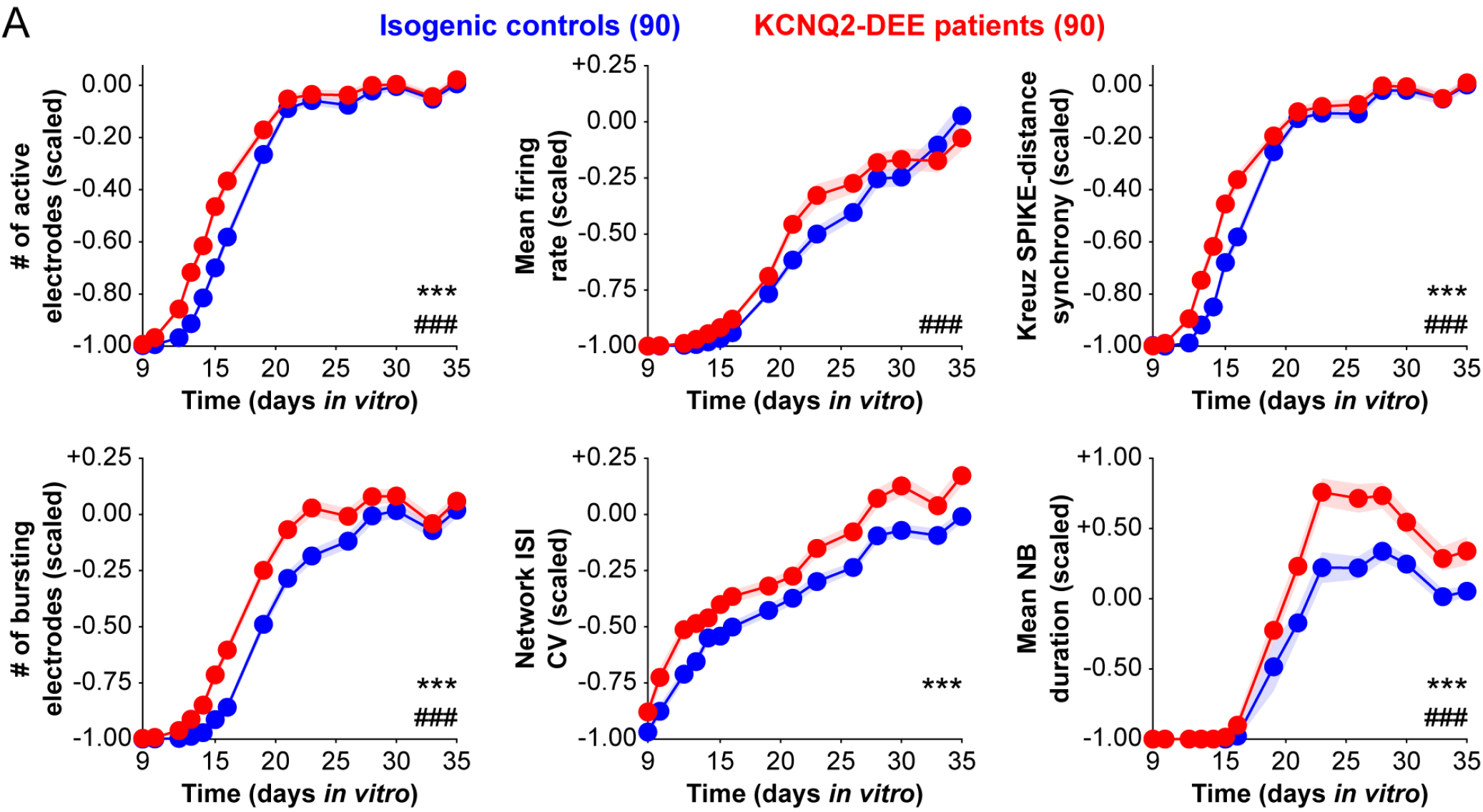
Longitudinal analysis of additional functional biomarkers associated with KCNQ2-DEE. **A)** Compared to pooled isogenic control neurons, pooled KCNQ2-DEE neurons exhibit an earlier onset of active and more synchronous neuronal populations as indicated by a higher number of active electrodes, increased mean firing rates, and elevated Kreuz SPIKE-distance synchrony. Additionally, pooled KCNQ2-DEE neurons show a greater number of bursting electrodes between weeks 2 and 4 in culture, prolonged mean network burst duration starting from week 3, and increased network ISI CV. Repeated-measures ANOVA: genotype effect: *p < 0.05, **p < 0.005, ***p < 0.0005; genotype/time interaction: # p < 0.05, ## p < 0.005, ### p < 0.0005. Data are shown as mean ± SEM, and the number of replicate wells is indicated in the panel.

**Figure S9.**
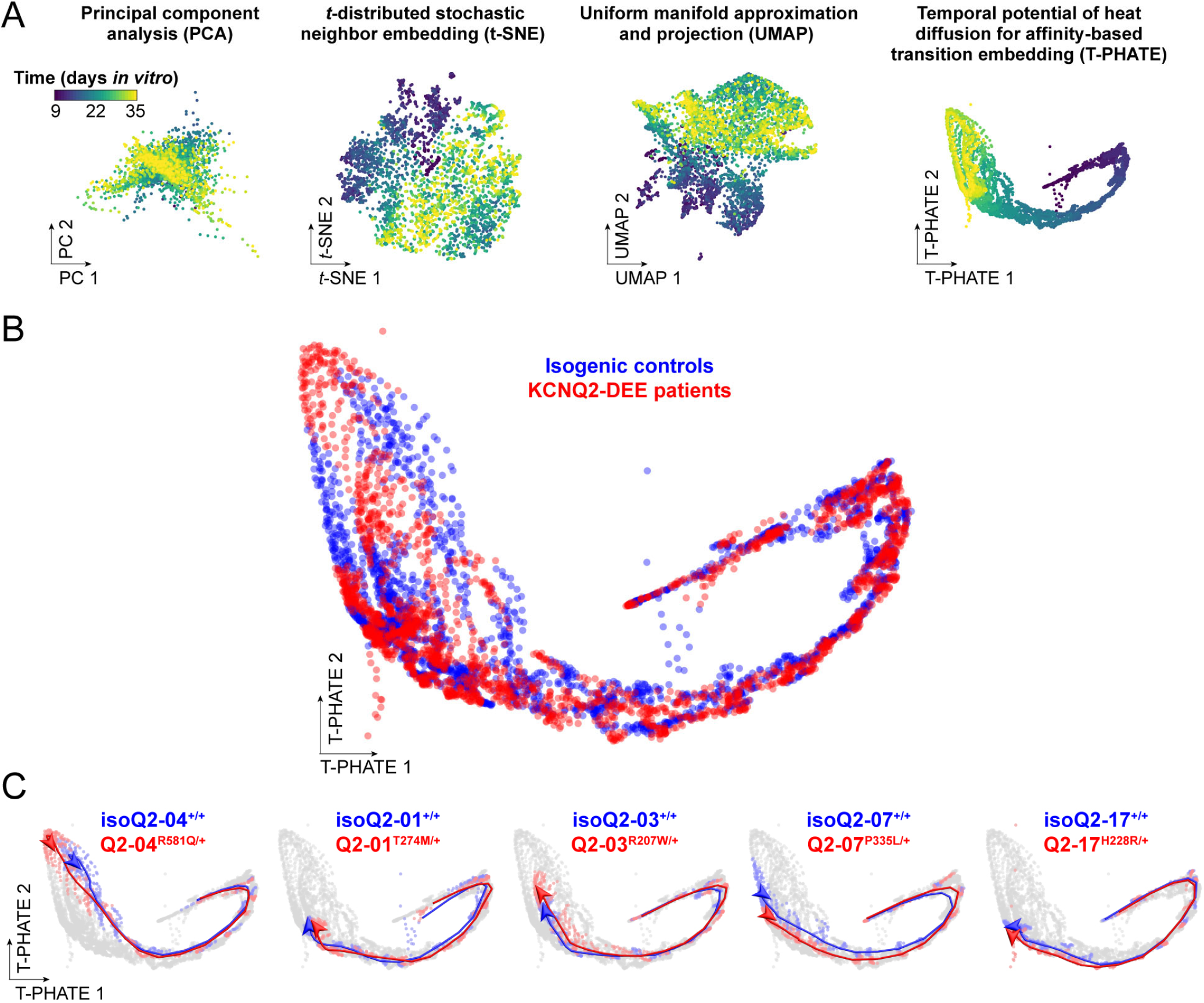
Functional trajectories of patient-specific KCNQ2-DEE iPSC neurons in low-dimensional spaces. **A)** Traditional dimension reduction techniques, such as principal component analysis (PCA), t-distributed stochastic neighbor embedding (t-SNE), and uniform manifold approximation and projection (UMAP) fail to preserve the temporal dynamics of neuronal development in MEA recordings. In contrast, temporal potential of heat diffusion for affinity-based transition embedding (T-PHATE), which integrates temporal autocorrelation, successfully captures the evolving dynamics of neuronal activity across time in a low-dimensional space. **B)** Enlarged overlay of all KCNQ2-DEE neurons and isogenic controls within the latent T-PHATE space from Figure 5C. **C)** Overlaying individual cell line labels onto the latent space demonstrates that functional variation is strongly structured by genetic background. Red (KCNQ2-DEE) and blue (isogenic control) lines indicate smoothed functional trajectories of individual cell lines.

**Figure S10.**
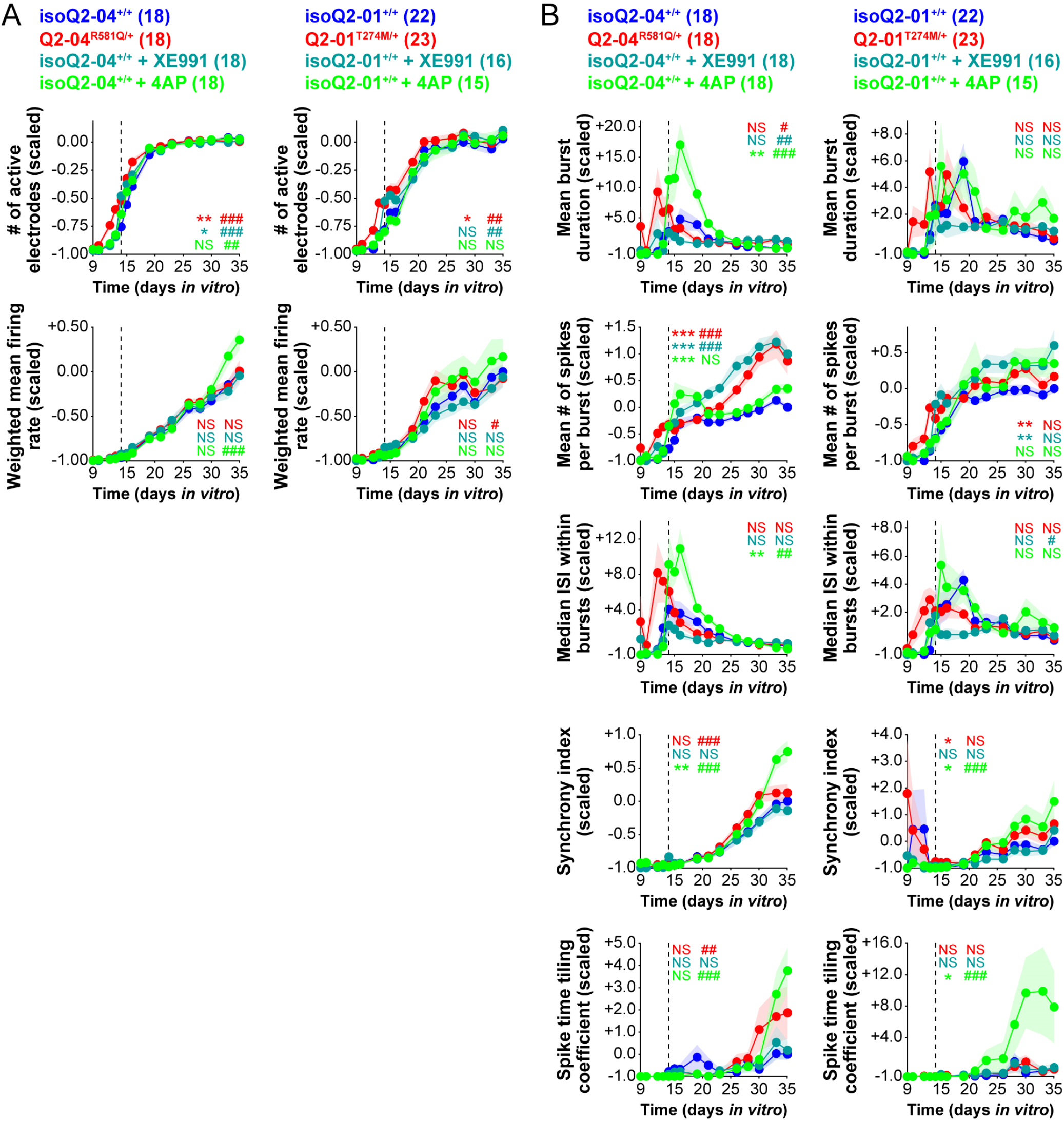
Machine learning predictions demonstrate that chronic KV7 inhibition selectively phenocopies KCNQ2-DEE functional signatures in isogenic control neurons, while inhibition of other K+ channels with 4AP leads to enhanced synchrony. **A)** Chronic XE991 significantly increased the number of active electrodes in both isoQ2-04+/+ (left) isoQ2-01+/+ (right) cultures while leaving the WMFR unchanged. In contrast, 4AP did not affect active electrode counts in either line but significantly elevated WMFR in isoQ2-04+/+ neurons, with no effect on WMFR in isoQ2-01+/+ neurons. Repeated-measures ANOVA on drug treatment days 14-35: genotype effect: *p < 0.05, **p < 0.005, ***p < 0.0005; genotype/time interaction: # p < 0.05, ## p < 0.005, ### p < 0.0005; NS: not significant. Colored symbols indicate significant differences between isogenic control and KCNQ2-DEE neurons (red), XE991-treated isogenic control (teal), and 4AP-treated isogenic control (green) neurons. Data are shown as mean ± SEM, and the number of replicate wells is indicated in the panel. **B)** Although, treatment with 4AP did not phenocopy KCNQ2-DEE functionally, it transiently increased the mean burst duration, mean number of spikes per burst, and median intra-burst ISI for several days after treatment onset in isoQ2-04+/+ neurons, with no significant effect in isoQ2-01+/+ neurons. At later time points both isoQ2-04+/+ (left) and isoQ2-01+/+ (right) cultures exhibited higher synchrony index and spike-time tiling coefficient, indicating greater network synchrony and not altered with XE991 treatment.

**Figure S11.**
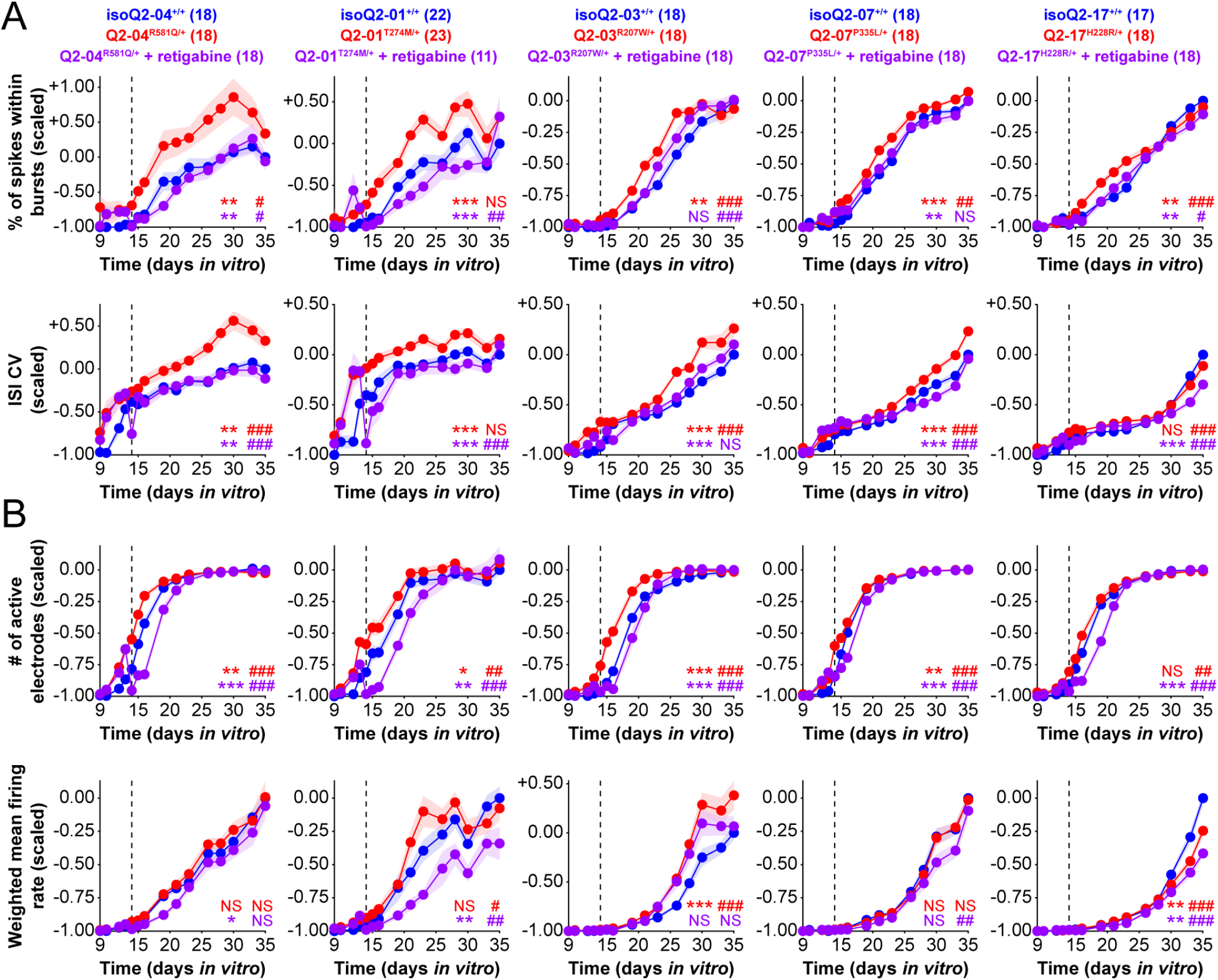
Machine learning predictions analysis demonstrates that the KV7 channel activator retigabine rescues KCNQ2-DEE phenotypes with variable efficacy. **A)** Chronic activation of KV7 channels with retigabine reduced irregular firing in KCNQ2-DEE neurons as indicated by decreased ISI CV and a reduced percentage of spikes occurring within bursts. Repeated-measures ANOVA on drug treatment days 14-35. genotype effect: *p < 0.05, **p < 0.005, ***p < 0.0005; genotype/time interaction: # p < 0.05, ## p < 0.005, ### p < 0.0005; NS: not significant. Colored symbols indicate significant differences between isogenic control and KCNQ2-DEE neurons (red), and between KCNQ2-DEE and retigabine-treated KCNQ2-DEE neurons (purple). Data are shown as mean ± SEM, and the number of replicate wells is indicated in the panel. **B)** Retigabine treatment significantly decreased the number of active electrodes in all 5 patient-specific KCNQ2-DEE iPSC neurons, and significantly reduced WMFR in Q2-04R581Q/+, Q2-01T274M/+, and Q2-17H228R/+ KCNQ2-DEE neurons, suggesting a dampening of overall spontaneous activity.

**Table S1.** Intrinsic membrane properties of KCNQ2-DEE patient and isogenic control neurons at days 22-26.

|  | Q2-01 |  | Q2-07 |  |
| --- | --- | --- | --- | --- |
| Genotype: | Q2-01<sup>T274M/+</sup> | isoQ2-01<sup>+/+</sup> | Q2-07<sup>P335L/+</sup> | isoQ2-07<sup>+/+</sup> |
| Resting Potential (mV) | -50.1 ± 1.5<br>(21) | -49.9 ± 1.8<br>(21) | -51.3 ± 1.1<br>(25) | -50.4 ± 1.4<br>(23) |
| Input Resistance R<sub>N</sub> at RMP (MΩ) | 890.5 ± 64.2 | 896.0 ± 69.3 | 785.0 ± 46.5 | 778.7 ± 51 |
| Series Resistance R<sub>S</sub> RMP (MΩ) | 12.0 ± 0.5 | 11.4 ± 0.4 | 11.0 ± 0.3 | 11.6 ± 0.3 |
| AP Amplitude from Baseline (mV) | 90.4 ± 2.2<br>(10) | 91.4 ± 2.1<br>(16) | 91.7 ± 1.4<br>(21) | 95.0 ± 1.8<br>(15) |
| AP Threshold (mV) | -30.3 ± 1.5 | -28.3 ± 1.5 | -28.0 ± 0.7 | -29.5 ± 1.1 |
| AP Half-Width (ms) | 2.3 ± 0.2 | 2.4 ± 0.4 | 2.4 ± 0.2 | 2.0 ± 0.2 |
| fAHP (mV) | -49.3 ± 1.8 | -48.6 ± 1.3 | -50.2 ± 1.0 | -51.6 ± 1.4 |
| mAHP (mV) | -4.7 ± 0.4<br>(21) | -4.0 ± 0.4<br>(20) | -4.1 ± 0.3<br>(24) | -3.7 ± 0.4<br>(21) |
| sAHP (mV) | -3.0 ± 0.3 * | -1.7 ± 0.4 | -2.2 ± 0.3 * | -1.2 ± 0.3 |

() indicate number of cells

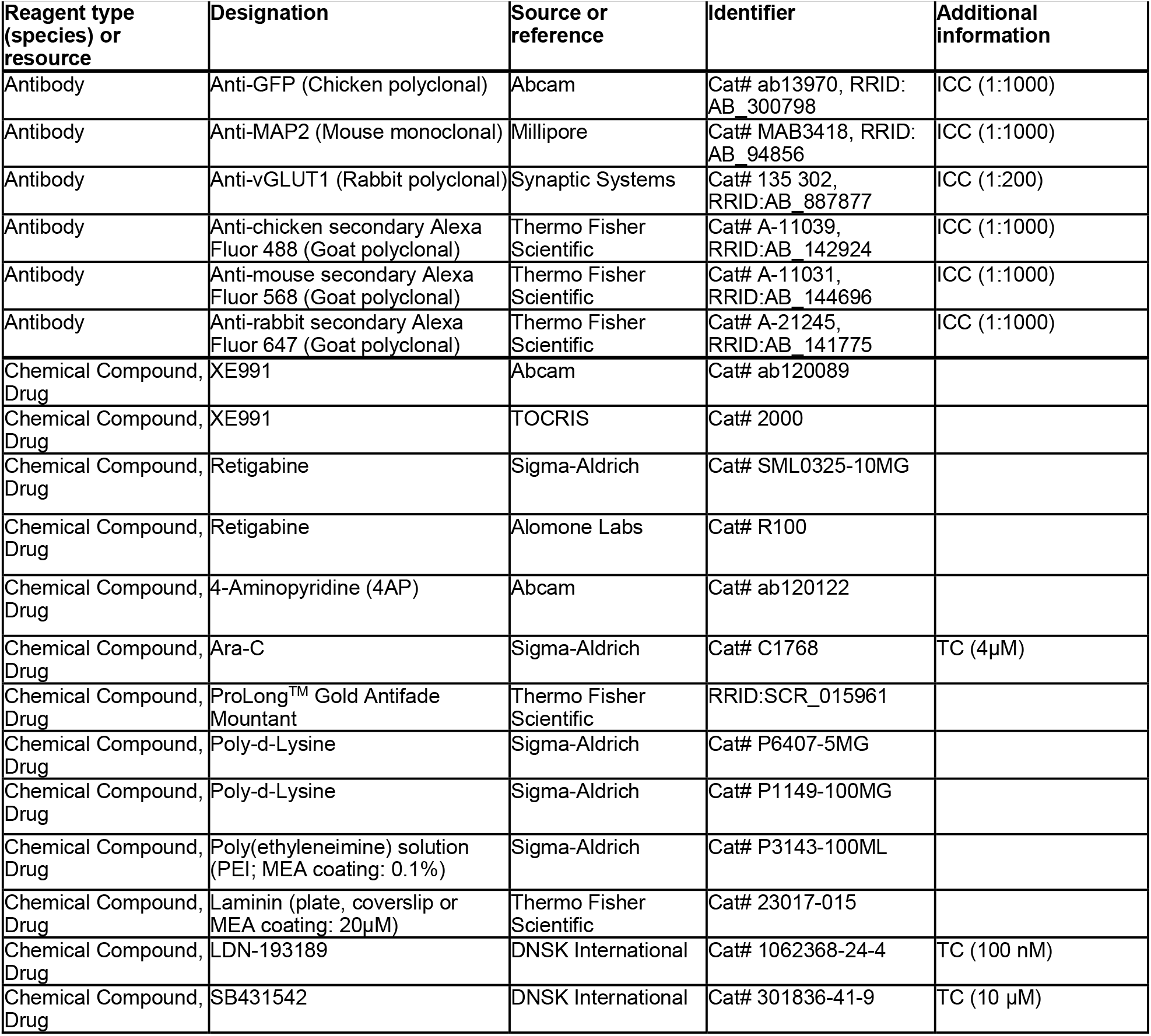
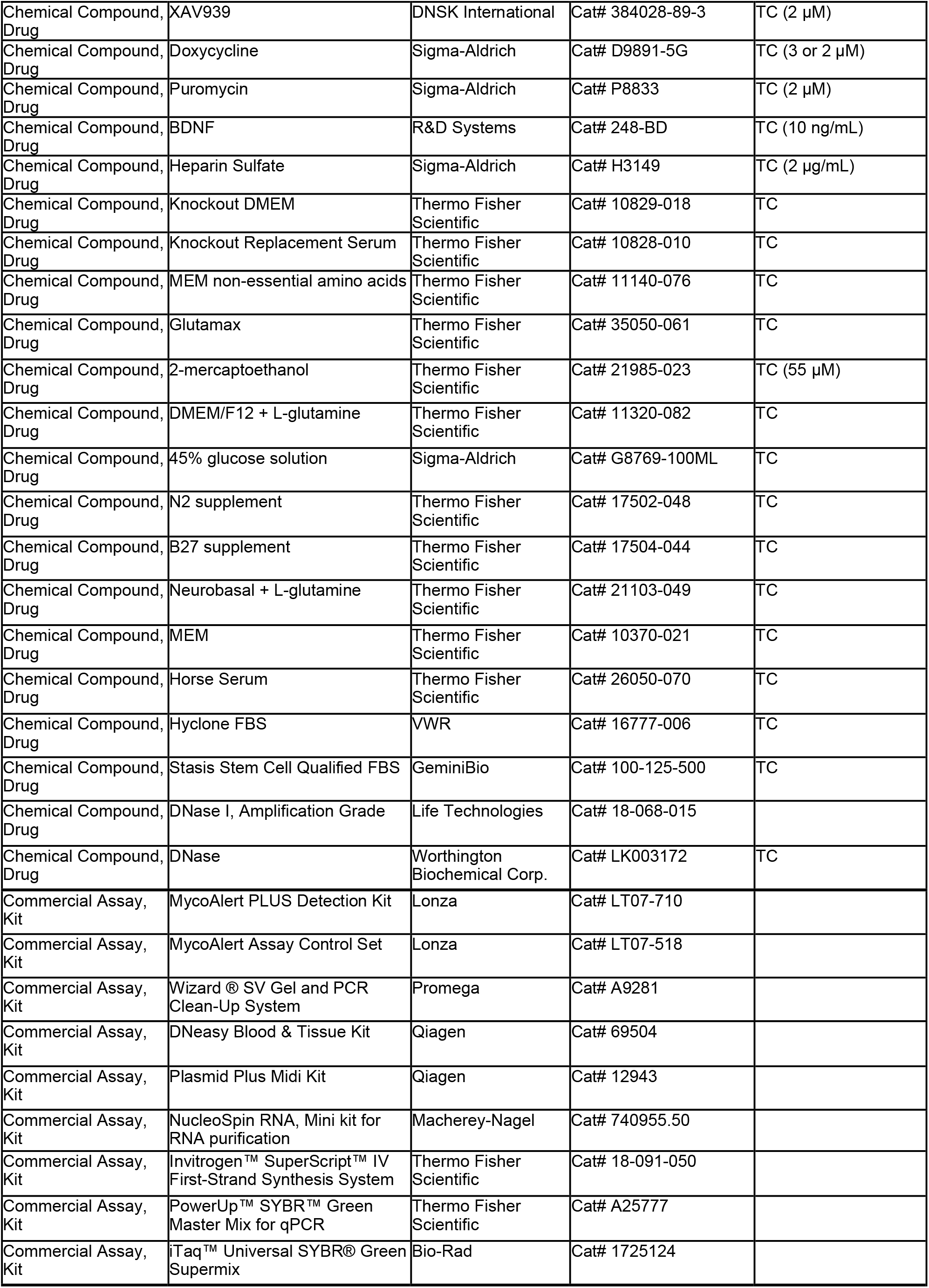
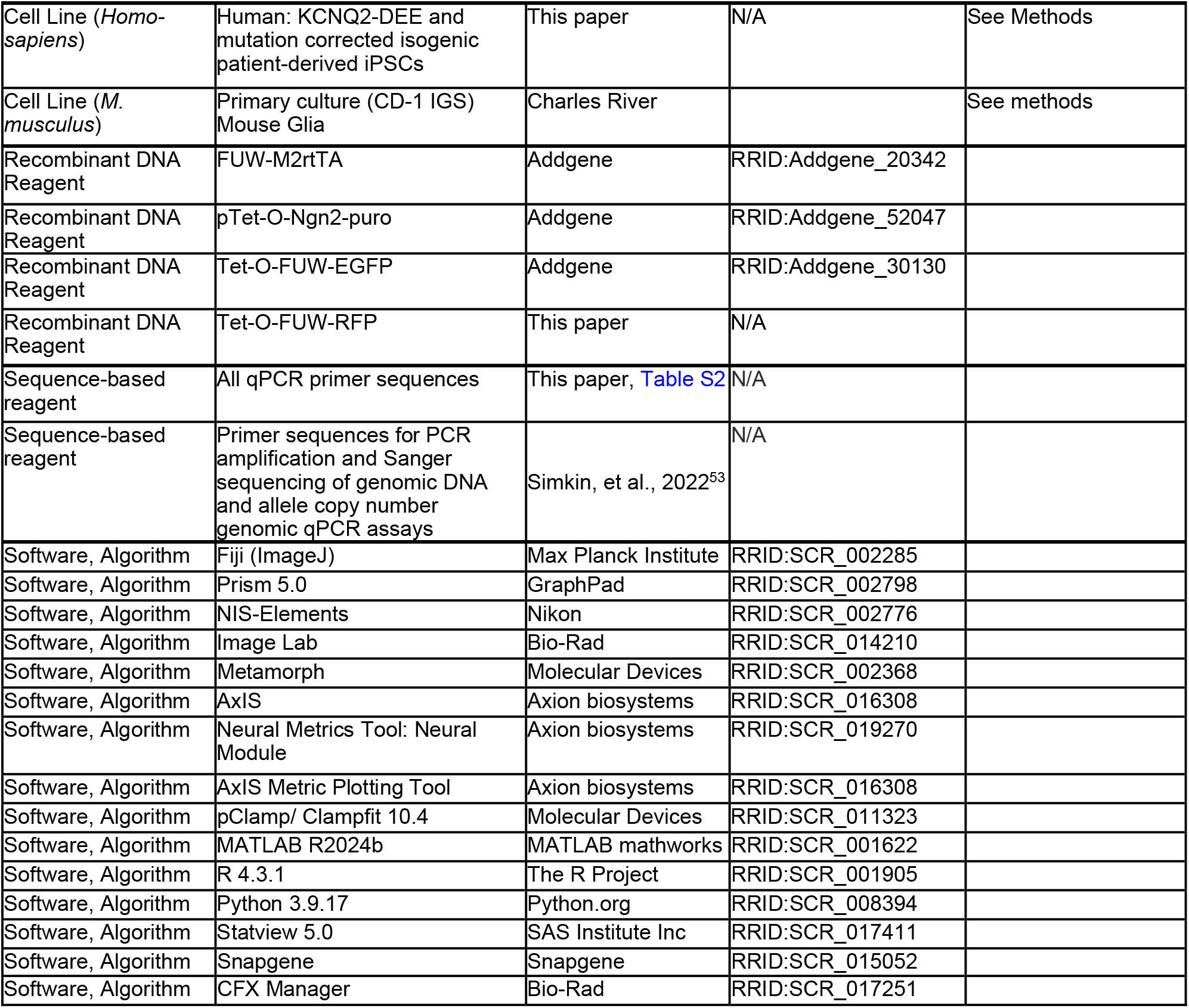
Key Resource Table.

